# Orco regulates the circadian activity of pheromone-sensitive olfactory receptor neurons in hawkmoths

**DOI:** 10.1101/2025.06.17.659282

**Authors:** Aditi Vijayan, Mauro Forlino, Yajun Chang, Pablo Rojas, Katrin Schröder, Anna C. Schneider, Martin E. Garcia, Monika Stengl

## Abstract

1

The mating behavior of nocturnal *Manduca sexta* hawkmoths is under strict temporal control. It is orchestrated via circadian and ultradian oscillations in sex-pheromone stimuli as social Zeitgeber. The extremely sensitive pheromone-detecting olfactory receptor neurons (ORNs) that innervate the long trichoid sensilla on the male’s antennae are peripheral circadian clocks. They express the transcriptional-translational feedback loop (TTFL) circadian clockwork, best characterized in *Drosophila melanogaster*. In hawkmoths, it is still unknown whether or how the ORN TTFL clockwork regulates the daily rhythms in pheromone sensitivity and in temporal resolution of ultradian pheromone pulses as prerequisites to the temporal regulation of hawkmoth mating behavior.

We hypothesize that, rather than the slow TTFL clock, a more rapidly adaptive post-translational feedback loop (PTFL) clockwork, assembled in a signalosome in the ORN plasma membrane, allows for temporal control of pheromone detection via generation of multiscale endogenous membrane potential oscillations. The potential oscillations of the PTFL clock could rapidly synchronize to oscillations of pheromone stimuli at different time scales, thus enabling the prediction of stimulus patterns as a mechanism for active sensing. With *in vivo* long-term tip recordings of long trichoid sensilla of male hawkmoths, we analyzed the spontaneous spiking activity indicative of the ORNs’ endogenous membrane potential oscillations. Consistent with our hypothesis of a multiscale PTFL clock in hawkmoth ORNs, spontaneous spiking was modulated on ultradian and circadian time scales, with maximum activity at night. When we blocked the evolutionarily conserved olfactory receptor coreceptor (Orco), the circadian modulation was abolished but the ultradian frequency modulation of the spontaneous activity remained. Consistent with PTFL control, Orco was not under the transcriptional control of the TTFL clock, but its modulation of spontaneous spiking activity was dependent on cAMP. We could replicate the experimental data in a conductance-based computational model of an ORN. In this model, Orco conductance changed as a function of fluctuating 2^nd^ messenger levels. This study demonstrates that a PTFL clock is sufficient to impose a circadian pattern on ORN sensitivity. Future experiments are necessary to further challenge our novel hypothesis.

**Significance statement:** It is generally assumed that all circadian rhythms in an organism are driven by a transcriptional-translational feedback loop (TTFL) clock. In this study, we demonstrate with *in vivo* recordings of hawkmoth pheromone-sensitive olfactory receptor neurons (ORNs) that the olfactory receptor coreceptor (Orco) is the key pacemaker channel for controlling circadian rhythms in spontaneous spiking activity. Since Orco expression is not driven by the TTFL clock, we put forward the novel hypothesis that Orco is a core element of a post-translational feedback loop (PTFL) membrane clock assembled in a signalosome controlling interlinked oscillations in 2^nd^ messengers and membrane potential. Accordingly, our computational model predicts that ORN sensitivity is tuned by circadian changes in the conductance of an Orco ion channel, which is mediated by cycling levels of cyclic nucleotides. Our novel hypothesis that highlights the contribution of feedback-controlled post-translational modifications to the generation of endogenous circadian rhythmicity should spark further challenges.

## 3 Introduction

Biological clocks generate endogenous physiological and behavioral oscillations on multiple timescales that persist in the absence of any external timing cues under constant conditions (Lamont and Amir, 2010). Such clocks are essential to generate internal rhythms and to permit their synchronization to external Zeitgebers at time scales from milliseconds to years. These synchronized endogenous multiscale clocks thus provide both a continuous time frame for physiology and behavior - the organism’s “present time” - and enable organisms to predict and respond to regular, periodic environmental changes (Stengl and Schneider, 2024). Even though endogenous clocks are widespread in prokaryotes and eukaryotes (Géron et al., 2024; Hall and Rosbash, 1993; Millar, 2016; Thaiss et al., 2014), current knowledge about how multiscale clocks are linked over widely diverging time scales remains sparse.

Insect circadian clock neurons, which are tightly connected to visual brain circuits, are amongst the best studied biological clocks (Hall and Rosbash, 1993; Nishiitsutsuji-Uwo and Pittendrigh, 1968; Reinhard et al., 2024). They regulate physiological and behavioral oscillations that underlie sleep-wake rhythms tied to the 24 h light-dark (LD) cycles on Earth (Dubowy and Sehgal, 2017). Circadian clock neurons possess a molecular clockwork of transcriptional-translational feedback loops (TTFLs). Entrained to the LD cycle, this TTFL clock generates endogenous oscillations of clock gene mRNA and protein levels with a cycle period of ∼24 h via negative feedback by proteins that inhibit their own transcription (Hardin, 2011). It is assumed for both insects and mammals that this TTFL comprises the master clockwork that regulates all physiological and behavioral rhythms in the circadian time range through genetically controlled output pathways (Hastings et al., 2020).

Molecular TTFL clockworks were not only found in central brain clock neurons but also in sensory neurons, such as olfactory receptor neurons (ORNs) in insect antennae, where they play a role in odor-dependent circadian behaviors like feeding, pollination, reproduction, and navigation (Flecke et al., 2010; Flecke and Stengl, 2009; Krishnan et al., 1999; Merlin et al., 2009, 2006; Page and Koelling, 2003; Plautz et al., 1997; Sauman and Reppert, 1998; Schendzielorz et al., 2015; Schuckel et al., 2007; Tanoue et al., 2004; Zhou et al., 2005). Little is known about how the TTFL clockwork of insect ORNs regulates circadian rhythms in chemosensory sensitivity that underlie the circadian control of these different behaviors.

At night, male nocturnal *Manduca sexta* hawkmoths are very sensitive to the sex pheromone blend that their conspecific females release in an intermittent pattern (Itagaki and Conner, 1988). The intermittency is a prerequisite to maintain the male’s search flight, which is suspended at pulse frequencies >30 Hz (Baker et al., 1985; Baker and Vogt, 1988; Stengl, 2010). The pheromone pulses are resolved of up to 3 Hz by pheromone-specific ORNs in the antenna’s long trichoid sensilla (Lee and Strausfeld, 1990; Marion-Poll and Tobin, 1992). Pheromone-specificity is provided by odor receptor (OR) subunits in the ciliary membranes of ORNs that heteromerize with the conserved olfactory receptor co-receptor Orco (Nakagawa and Vosshall, 2009; Stengl and Funk, 2013; Wicher and Miazzi, 2021). The insect odor transduction cascade is still under debate and seems to involve both ionotropic and metabotropic pathways (Sato et al., 2008; Schneider et al., 2025; Wicher et al., 2008). While there is no general agreement on all roles of Orco, it is known that in heterologous expression systems Orco homo- and heteromers form a slow, leaky, non-specific cation channel, which continuously depolarizes the cells (Jones et al., 2011; Sargsyan et al., 2011; Sato et al., 2008; Wicher, 2018; Wicher et al., 2008). *In vivo* tip recordings of long trichoid sensilla in *M. sexta* confirmed that Orco acts as leaky cation channel, acting as a pacemaker channel, because it promotes ultradian membrane potential oscillations that underlie the spontaneous action potential activity (Nolte et al., 2013). These potential oscillations turn the ORNs into ultradian temporal filters that are sensitive to the frequency of the pheromone pulses they encounter. Pheromone pulses that arrive at a similar frequency as the endogenous sub-threshold membrane potential oscillations would synchronize these oscillations and would be more likely to elicit action potential trains. Therefore, this pacemaker channel-dependent mechanism would allow for active sensing if endogenous membrane potential oscillations are in the physiological range that is necessary for tracking intermittent pheromone pulses.

It is not known whether the Orco-dependent membrane potential oscillations also occur at circadian time scales, and whether or how they are linked to ultradian potential oscillations. One hypothesis is that the molecular TTFL clockwork of ORNs exclusively controls endogenous circadian rhythms of the membrane potential via circadian expression of pacemaker channels like Orco. As alternative hypothesis, the plasma membrane of ORNs could by itself comprise a signalosome that constitutes an autonomous clock that is entrained by, but does not rely on, the TTFL clock. Constant levels of colocalized ion channels with antagonistic effects that do not require daily degradation and daily transcription via the TTFL clock, but with modulation of conductance by post-translational modifications, would constitute a more economical and faster adjustable mechanism to generate endogenous membrane potential oscillations (Stengl and Schneider, 2024). This would facilitate active sensing by signalosome-anchored receptors, rapidly tuning odorant/pheromone detection to behaviorally relevant Zeitgeber signals in the insect’s environment, other than the LD cycle that is linked to the TTFL clock.

Here, we performed a combination of experimental and computational modeling studies to investigate whether Orco plays a role in active sensing at the circadian and/or ultradian time scale via generation of endogenous membrane potential oscillations. We hypothesize that Orco is part of a post-translational feedback loop (PTFL) clockwork, assembled in a signalosome in the ORN somatic plasma membrane, rather than being TTFL controlled. To capture naturally occurring, physiologically relevant processes in insect ORNs we performed *in vivo* long-term tip recordings of the spontaneous spiking activity of pheromone-sensitive trichoid sensilla of intact but restrained hawkmoth males.

We corroborated the experimental data with a Hodgkin-Huxley (HH) based computational ORN model. In general, the widely used deterministic HH models do not include circadian control of ion channels. Moreover, the highly dynamic, bursting firing patterns of ORNs cannot be described by the traditional deterministic modeling of neuronal activity. This constitutes a drawback because stochasticity plays a crucial role in shaping firing patterns which are influenced not only by external stimuli but also by the inherent noise involved in channel gating. Therefore, to address the above points realistically, we developed a Langevin formulation of the HH model, which explicitly accounts for the noise introduced by the random opening and closing of ion channels and circadian modulation of the Orco channel conductance. By integrating channel noise, we could explore how circadian regulation of ionic currents contributes to the variability of firing patterns and how this noise interacts with external stimuli to influence olfactory signal processing.

Our model aims to highlight the advantages of using a stochastic framework for modeling the circadian activity of ORNs. The combined experimental and modeling efforts generate a more complete picture and predictions concerning these autonomous processes in sensory neurons, which are required for future exploration of physiologically relevant anticipatory behaviors that are driven by biological clocks.

## 4 Methods

### 4.1 Animals

Nocturnal *M. sexta* hawkmoths were bred and raised from eggs in the rearing facility at the University of Kassel. Males and females were separated during pupal stages. Males were housed isolated from the females to avoid exposure to female pheromones. All experiments were performed with adult males reared in a long-day photoperiod (L:D 17:7 h to prevent diapause) at 25 °C with relative humidity of about 55%. Larvae were fed with an artificial diet modified after Bell and Joachim (1976); adult moths could feed on sugar solution with added vitamins (Roth) *ad libitum* (Riffell et al., 2008).

While all animals were exposed to the same lighting regime as Zeitgeber to entrain them up until the start of the experiments, we also attempted pheromone exposure as additional circadian Zeitgeber to phase-align spontaneous ORN activity in a subset of males (Ghosh et al., 2024). Without allowing direct contact, virgin males were exposed at Zeitgeber time (ZT) 16 to females and naturally occurring pheromone blends in closed mating cages for 30 mins. Males were then moved back into isolation before tip recordings (see below) were started at ZT 17 on the same day. Since we focused on spontaneous spiking activity rather than on pheromone-dependent responses and because repetitive or long pheromone exposure changes the composition of 2^nd^ messenger modulated ion channels in ORNs (Gawalek and Stengl, 2018), we did not employ any further pheromone synchronization protocols.

### 4.2 Solutions

Ringer compositions were taken from (Pézier et al., 2007) and contained (in mM): 6.4 KCl, 12 MgCl_2_, 1 CaCl_2_, 12 NaCl, 10 HEPES, 340 glucose for hemolymph ringer (HLR) at 450 mosmol/kg, and 172 KCl, 3 MgCl_2_, 1 CaCl_2_, 25 NaCl, 10 HEPES, 22.5 glucose for sensillum lymph ringer (SLR) at 475 mosmol/kg, both at pH 6.5.

The Orco antagonist OLC15 (N-(4-butylphenyl)-2-((4-ethyl-5-(2-pyridinyl)-4H-1,2,4-triazol-3-yl)thio)acetamide (Chen and Luetje, 2012); kindly provided by Dr. D. Wicher, Max Planck Institute for Chemical Ecology, Jena, Germany) was dissolved in DMSO to a concentration of 100 mM (stock solution stored at 4-8 °C) before being diluted in SLR to a final concentration of 50 µM to be used within one week (stored at 4 °C). We determined in previous studies that, compared to other Orco antagonists, OLC15 is the most specific (Chen and Luetje, 2012; Nolte et al., 2016, 2013; Pask et al., 2013). It is effective at 50 µM, according to dose-response curves, and dose-dependently antagonizes the action of the Orco agonist VUAA1 in intact hawkmoths (Nolte et al., 2016, 2013). DMSO percentage was 0.05% in the final solution.

The cAMP analog 8-bromo-cAMP acetoxymethyl ester (8-Br-cAMP-AM; Biolog Life Science Institute GmbH & Co. KG) was dissolved in DMSO to a concentration of 10 mM (stock solution stored at −20°C) and diluted to a final concentration of 100 µM in either SLR or SLR + 50 µM OLC15.

### 4.3 Electrophysiology

The spontaneous electrical activity of the unstimulated pheromone-sensitive ORNs in a single long trichoid sensillum was recorded extracellularly as *in vivo* tip recording (Kaissling et al., 1987). 1-2-d-old male *M. sexta* that were raised and entrained to Zeitgebers as described above were fixed in a custom-made Teflon holder with adhesive tape. The right antenna used for recordings was oriented with its dorsal surface facing up and immobilized near the base with dental wax (Boxing wax strips, KerrHawe SA). Electrodes (Ag/AgCl wire in a glass pipette filled with SLR for the recording electrode and HLR for the reference electrode) were pulled (DMZ-Universal Puller, Zeitz Instruments) from borosilicate capillaries (OD:1.56 mm, ID: 1.17 mm, without filament; Science Products) to a tip diameter of about 2 μm. All recordings were made at a distance of 2/3 to 3/4 of the antenna length towards its distal end because pheromone-sensitive long-trichoid sensilla (sensilla trichodea type 1) occur in a characteristic, homogenous pattern along the hawkmoth antenna (Kaissling et al., 1989; Kalinová et al., 2001; Lee and Strausfeld, 1990). Furthermore, all ORNs in long trichoid sensilla express the same Orco staining pattern (Nolte et al., 2016), share largely overlapping ion channel populations, and obvious differences in the activity of ORNs from different annuli have not been reported (c.f., Dolzer et al., 2003; Gawalek and Stengl, 2018; Schneider et al., 2025). Thus, it is very likely that pheromone encoding mechanisms are not locally restricted along the flagellum. Recording and reference electrodes were located on the same annulus: After cutting off the distal 20 annuli of the antenna with micro scissors, the reference electrode was inserted to a depth of 2 annuli into the antennal lumen from the cut end. The cut was covered with electrode gel (GE Medical Systems Information Technology) to prevent desiccation. Under microscopic control, some tips of the long trichoid sensilla of the annulus were cut off with sharpened Dumont forceps, and the recording electrode was slipped over a single sensillum of an upper sensillar row. Electrodes were moved with manual micromanipulators (MMJ, Märzhäuser Wetzlar; and Leitz, Leica Microsystems) or with motorized micromanipulators (Microstar, Scientifica) with a joystick controller (Scientifica).

ORN activity was amplified 200-fold (custom built amplifier with 10^12^ Ω input impedance, or ELC-01 MX in combination with DPA-2FX, npi, or EXT 10-2F, npi), low pass filtered at 1.3 kHz, digitized at 20 kHz with a Digidata (1550A or 1550B, Molecular Devices), and saved for offline analysis with Clampex 10 (pCLAMP suite, Molecular Devices).

All tip recording experiments were performed at room temperature (21-27 °C) and a relative humidity of 35-60%. These environmental fluctuations across experiments were based on seasonal influences. Throughout each experiment, conditions were stable. To ensure that neither electrode reattachment to the same trichoid sensillum nor application of DMSO influenced spiking activity, we performed two sets of paired experiments at ZT 1-3 in which we recorded the spontaneous activity before and after removing and reattaching the SLR-filled recording electrode to the same sensillum, or recorded the spontaneous activity of the same sensillum in SLR and SLR + 0.15% DMSO. Both treatments did not affect spontaneous spiking activity (Figure 1B).

**Figure 1:**
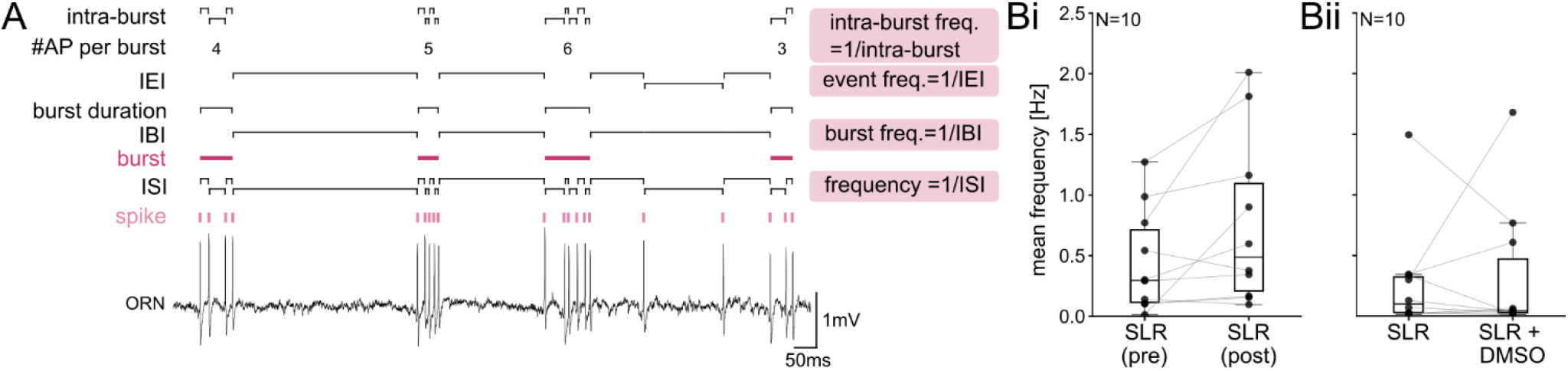
Attributes of pheromone-sensitive ORN spiking patterns. **(A)** Spikes were identified by their prominence in the raw ORN recording. The inter-spike interval (ISI) was defined as the time from one spike to the next. A train with ≥2 spikes with ISI ≤ 50 ms was considered a burst. The inter-burst interval (IBI) was the ISI between the last spike of the first burst and the first spike of the subsequent burst, omitting any individual spikes in between. Burst duration was the time between the first and last spike of a burst. The inter-event interval (IEI) included the ISI between bursts and individual spikes. The number of action potentials (spikes) per burst, and percentage of total action potentials as part of bursts were computed. The intra-burst interval was the ISI within a burst. Frequency of spikes, bursts, events and intra-bursts were determined as inverse of ISI, IBI, IEI and intra-burst interval. Detaching and reattaching the recording electrode to a long trichoid sensillum (**Bi**; paired t-test t(9) = −2.184; p = 0.057; N = 10) or adding DMSO to the sensillum lymph ringer (SLR) (**Bii**; Wilcoxon signed-rank test Z = 0.153; p = 0.922; N = 10) did not affect the mean spiking frequency.

We recorded ORN activity either in long-day conditions (17:7 LD) or constant darkness (DD) to examine the free-running peripheral ORN clock. For LD experiments, the light regime continued as in the rearing facility during the experiment. For DD and OLC15 experiments, the lights were turned off at the beginning of the recording, i.e., all animals were exposed to the same light Zeitgeber until the beginning of the experiment. To examine the effect of Orco on the spontaneous action potential (AP) activity pattern, we used the recording electrode to infuse OLC15 in DD conditions. To examine the effect of cAMP on Orco gating, we recorded the spontaneous ORN activity at the end of the moth’s activity phase (ZT 1-3) in SLR as control, and either 8-Br-cAMP-AM, or 8-Br-cAMP-AM + OLC15, infused through the recording electrode.

ORNs generate spontaneous action potentials. However, high spiking activity during control recordings is indicative of stress, whereas low/no activity could be indicative of damage. Therefore, we only considered experiments in which the average spiking frequency was between 2 Hz and 0.008 Hz in control conditions.

### 4.4 Data analysis

To characterize the spontaneous ORN spiking activity, we first detected spikes with a self-written program in Python. Using the scipy.signal package and find_peaks function, we used a moving mean with 100 samples (5 ms) window size for baseline correction, followed by applying a 8^th^ order low-pass Bessel filter (2000 Hz cutoff frequency) to the raw voltage trace to facilitate spike detection within the following conditions: spike full width (3-15 ms), threshold search for amplitude of spike (0.5-2.5 mV) and threshold prominence (0.1). Two or more consecutive spikes with intervals ≤ 50 ms were considered a burst (Dolzer et al., 2001). We then calculated the following attributes in bins of either 10 min or 1 h for further analysis: instantaneous spike frequency (ISI, 1 / inter-spike interval), inter-burst interval (IBI, time between two consecutive bursts with no single spikes in between), inter-event interval (IEI, time between two events, i.e. two consecutive single spikes, two consecutive bursts, a burst and a consecutive spike, and vice versa), burst duration (time from first to last spike in a burst), percentage of all spikes in the experiment that are members of burst, and mean number of spikes per burst (Figure 1A). All statistics were done with those binned values.

Hawkmoth behavioral and physiological rhythms desynchronize across the population in the absence of Zeitgebers like circadian rhythms of pheromone stimulation and daily light-dark cycles. Therefore, we first used RAIN analysis (Thaben and Westermark, 2014) to identify circadian rhythms (24 ± 4 h) in the attributes above for each animal and condition (LD, DD, OLC15) in the 1 h-binned data. Data was downsampled to 2 h resolution and thus included every other 1 h bin to speed up computational time. To further quantify the circadian changes in attributes across animals, it was then necessary to phase-align the recordings obtained in DD and OLC15. To achieve this, we used an optimization procedure to calculate the optimal temporal lags for each dataset, aiming to maximize the overall cross-correlation between the calculated binned spike frequency series. The optimization was performed using simulated annealing, which provided a set of time shifts (lags) for each series. These lags were applied to shift the frequency data in a manner that maximized the synchronization between the different subjects’ data, resulting in a “subjective ZT” with subjective ZT 0 at the first maximum of the binned ISI for each individual. The objective function used for optimization was the negative sum of pairwise correlations between all series after applying the shifts. The resulting alignment allowed us to compare the frequency patterns across the different conditions.

Once the data were aligned, we computed the mean frequency across all subjects for each bin and each condition. Then, the first high activity period was identified by finding the maximum in the mean frequency series. Subsequently, we identified the second high activity period as the time 24 hours after the first one and the first low activity period was found by locating the minimum in the mean frequency series between the two high activity periods. Besides the mean frequency, we also searched for significant oscillations in IBI, IEI, burst duration, percentage of spikes that are part of bursts, and mean number of spikes per burst. For this we found the maximum value of these statistics within a 3 h range centered in the first low activity period and compared it with the values at the second high activity period.

Continuous wavelet and Fourier analysis were performed using PyBoat (version 0.9.12) (Mönke et al., 2020) to examine the time series of neuronal firing frequency. The continuous wavelet method allows us to detect and characterize rhythmic patterns within the data by identifying frequency components that change over time. Specifically, we used wavelet analysis to determine whether a neuron exhibited circadian and ultradian rhythms. Unlike traditional Fourier analysis, which provides an overall frequency composition but assumes stationarity, wavelet analysis is particularly useful for biological signals because it can reveal how these frequencies evolve dynamically. By applying this approach, we could assess not only the presence of ultradian rhythms but also whether their strength or occurrence fluctuated in a circadian manner, indicating modulation by the 24-hour cycle. For this analysis, we used the 10 min-binned data to have sufficiently high resolution to detect ultradian rhythms. Periods were determined as local maxima in the Fourier power spectra.

### 4.5 Real-time quantitative polymerase chain reaction (qPCR)

We followed the qPCR methodology detailed in (Schneider et al., 2025). Briefly, male moths were isolated and cultured until the second day after eclosion, and the antennae from four individuals were collected every 4 hours for one day, starting at ZT 0, using liquid nitrogen, with four biological replicates per time point. Total RNA was extracted using TRIzol® Reagent (Invitrogen) and quantified for concentration and purity with the NanoDrop ND-1000 spectrophotometer (Thermo Fisher Scientific). First-strand complementary DNA (cDNA) was synthesized using the PrimeScript™ RT reagent Kit with gDNA Erase (Takara Bio Inc.). Primers for *Orco* (F: GCTCGCTACCACCAAATTGC, R:TCGTGACCCAACTGACAACA; Gene ID: LOC115452348) and for *timeless* (*tim*) (F: TTAAGCCGACCGTAGTGCTG, R: CGTCTTCCGTCCATGTGTCT; Gene ID: LOC115446009) were designed using the Primer-BLAST online program of NCBI. The qPCR was conducted following the protocol of TB Green® Premix Ex Taq™ II (Tli RNase H Plus) (Takara Bio Inc.), with the program: 95 °C for 30 s, followed by 40 cycles of 95 °C for 5 s, and 60 °C for 30 s. Relative expression levels of *Orco* were calculated using the 2^-ΔΔCt^ method (Livak and Schmittgen, 2001).

### 4.6 Statistics

All statistics were done with self-written python codes, SigmaPlot 12 (Systat Software), or JASP (versions 0.19.03 and 0.95.4, University of Amsterdam) with a significance level α = 0.05. Unless noted otherwise, data were checked for normal distribution with Anderson-Darling or Shapiro-Wilk tests and for equal variance with Levene’s test. If one of these assumptions failed, we used non-parametric tests, otherwise we used parametric tests.

To assess the statistical significance of observed differences in neuronal activity of the aligned data between ZTs, we applied the Wilcoxon signed-rank test to pairwise comparisons between the first low activity period and the subsequent high activity period and discarded the transient high activity at the beginning of most recordings.

For paired recordings (electrode reattachment, addition of DMSO, or cAMP/cAMP+OLC15 in the same animal, we used either paired student-t test or Wilcoxon signed-rank test, depending on normal distribution of data.

For other data, we used an ANOVA on ranks with either Tukey’s or Dunn’s post-hoc test for multiple comparisons as indicated.

### 4.7 ORN Model

In this study, we introduce a single-compartment Hodgkin-Huxley (HH) type model of the spontaneous activity of *M. sexta* ORNs (available on GitHub (Forlino et al., 2025)). The model contains 5 sets of ion channels that were previously identified in electrophysiological recordings (Dolzer et al., 2021): the fast sodium (Na^+^) and potassium (K^+^) channels responsible for the generation of spikes, low voltage-activated calcium channels (LVA) responsible for the initiation of bursts, calcium-gated potassium channels (BK) responsible for the termination of bursts, and the Orco channel, which is a voltage-independent non-specific, cAMP-gated channel. A schematic of the model is provided in Figure 2A. We modeled Orco as a ZT-dependent conductance to represent the effect of the oscillations in cAMP concentration that were identified in both hawkmoth and cockroach antennae (Schendzielorz et al., 2014, 2015, 2012) (Figure 2B, C). For simplicity we chose to use a sine function to directly represent the oscillations in Orco conductance, but similar results can be achieved by modeling it as an output of a Goodwin model of the circadian clock via cAMP concentration.

**Figure 2:**
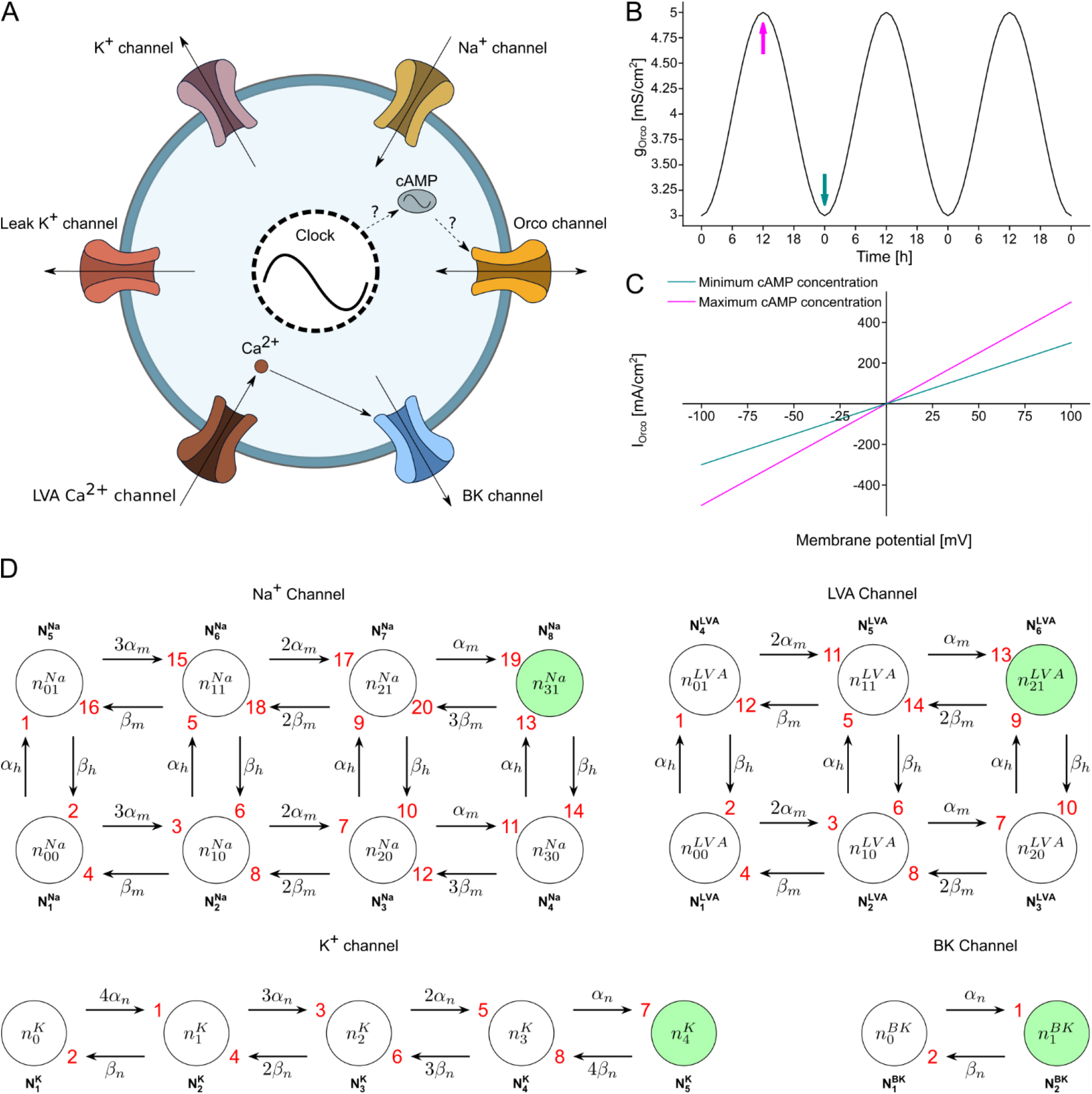
The conductance-based model of a simplified hawkmoth ORN. **(A)** Schematic of the model containing a voltage gated sodium channel (Na^+^), a voltage gated potassium channel (K^+^), a low voltage activated calcium channel (LVA), a calcium and voltage activated potassium channel (BK), a leak channel, and a cAMP-gated leaky non-specific cation channel (Orco). The Orco channel is represented as a linear conductance whose value oscillates with a sinusoidal shape as a function of the ZT. The circadian rhythm in the Orco conductance is hypothesized to result from a cAMP-dependent increase in channel open-time probability (as found in *Drosophila* Orco: (Getahun et al., 2013)) via a circadian oscillation of antennal cAMP concentrations (Schendzielorz et al., 2015). This oscillation is driven by the circadian clock and serves as an input; the internal mechanism of the clock is not modeled explicitly. Arrows indicate the flow of ions through the channels. **(B)** Parametrized circadian oscillation in Orco conductance introduced as an external input to our model, representing the gating of Orco by cAMP whose concentration oscillates on a circadian time scale. The times of minimum and maximum cAMP concentration are indicated by colored arrows. **(C)** Parametrized IV curve of the Orco channel showing a linear behavior with the slope depending on the cAMP concentration. Line colors correspond to the arrows in **B**. **(D)** Markov-chain representing all possible states and transitions for each type of ion channel included in the model. The per capita transition rates (α and β) depend on membrane potential and in the case of the BK channel also on the intracellular Ca^2+^-concentration. The “N” state vectors contain the population of channels in each state “n”. Directed edges are numbered in red. The only conducting state of each ion channel, representing the condition where all its gates are open, is shown in green.

Each type of ion channel possesses one or more gates that can open and close individually with transition rates α and β for opening and closing, respectively, of activation gates (n, m) and inactivation gates (h). As a result, each possible combination of open and closed gates defines a possible state in which each ion channel can be found. A schematic of the Markov chain depicting all possible states and transitions for each ion channel is shown in Figure 2D. Each ion channel can conduct current only in the state where all its gates are open.

First, we constructed a deterministic 22-dimensional model to serve as the mean field of the Langevin model following the methodology proposed by Pu and Thomas (2020). Here, one dimension tracks the membrane potential, another the intracellular Ca^2+^ concentration, and the remaining 20 dimensions track the number of channels that are in a specific state of the Markov chains. Considering the conservation of the total number of ion channels, each type of ion channel required one less dimension than the total possible states.

The complete Langevin model adds to the mean field an independent noise source for each transition in the Markov chain. These noise sources are biophysically motivated since the transition between ion channel states is a stochastic process, with the probability of transition determined by the kinetics of each kind of ion channel. The variability in the spiking behavior of a neuron emerges from this non-deterministic gating of individual ion channels. The detailed mathematical description of the model can be found in the Supplementary Material.

## 5 Results

We tested the hypothesis that sensory neurons, such as insect ORNs, perform active, anticipatory sensing based upon an endogenous plasma membrane clock with the pacemaker channel Orco as the core element. We focused on pheromone-sensitive long trichoid sensilla of male *M. sexta* hawkmoth antennae which express daily rhythms in pheromone sensitivity and temporal resolution that are controlled by a circadian clock. We performed minimally invasive *in vivo* tip recordings over the course of several days to search for daily rhythms in spontaneous spiking activity as a measure of endogenous membrane potential oscillations.

### 5.1 Pheromone-sensitive hawkmoth ORNs exhibit Orco-dependent circadian rhythms in spontaneous spiking activity

The ORNs that innervate the pheromone-sensitive long trichoid sensilla on male *M. sexta* antennae display spontaneous spiking patterns of irregular bursts with interspersed single action potentials in short-term recordings (Dolzer et al., 2001). In the present study we documented the same pattern in long-term recordings over the course of 2-7 days. This enabled us to search for daily and circadian rhythmicity in spontaneous ORN spiking activity in the absence of pheromone stimulation.

RAIN analysis of the spontaneous spiking patterns revealed significant daily rhythms in 7 of 11 animals under long-day conditions (17:7 h LD) (Figure 3A). The maxima in spontaneous activity occurred mostly during the dark phase when nocturnal hawkmoths are active (Figure 3A (top), Bi light blue arrow), and minima occurred during the light phase when hawkmoths rest/sleep (Figure 3A (bottom), Bi dark blue arrow). High spiking activity in the dark phase had a mean maximum frequency of ∼3 Hz (delta range: 0.5 – 4 Hz) (Figure 3Bii). In contrast, the minimum spiking frequencies during the hawkmoth’s resting phase were below 0.5 Hz (Figure 3A (bottom)).

**Figure 3:**
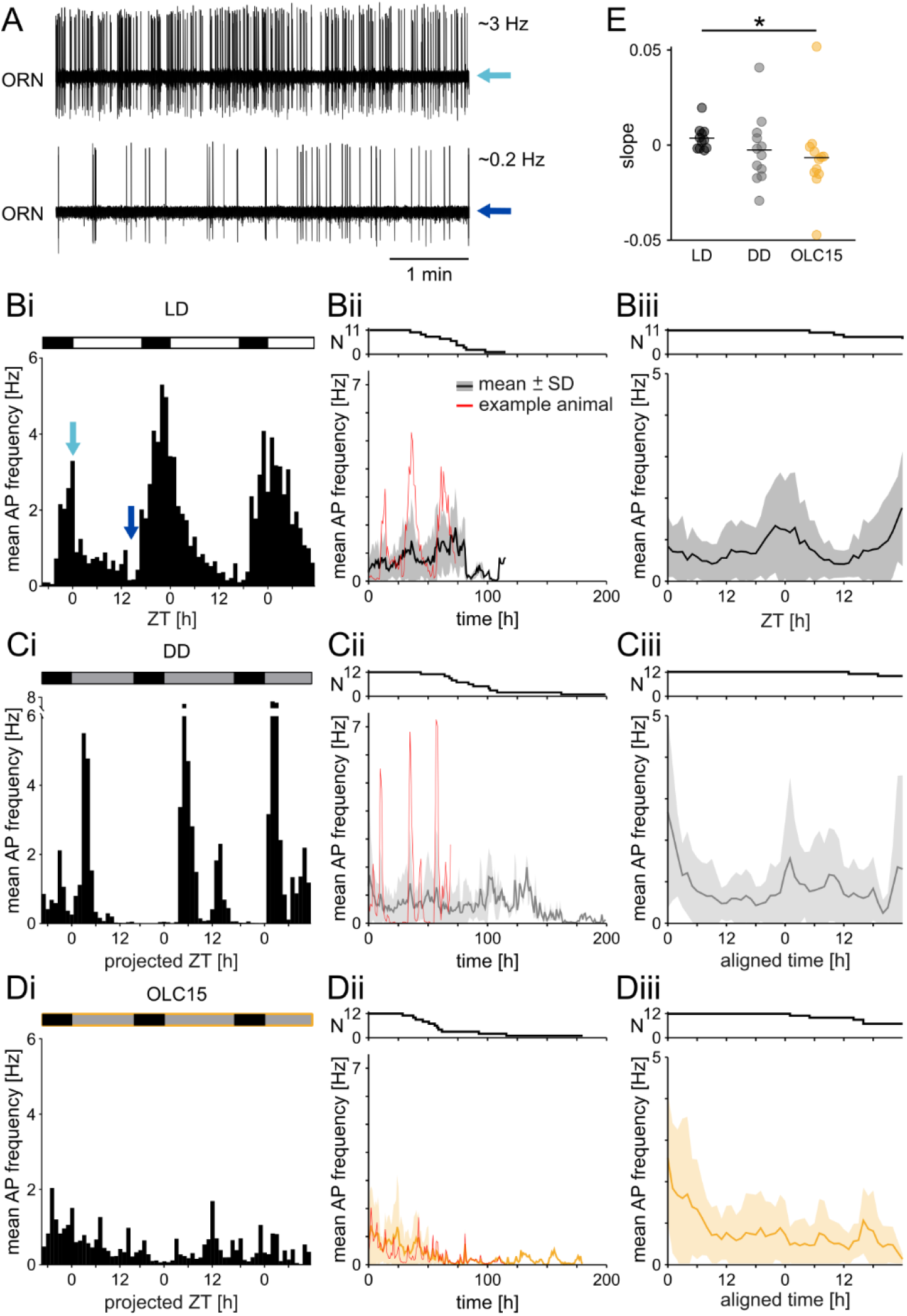
Spontaneous activity of pheromone-sensitive olfactory receptor neurons (ORNs) of male *M. sexta* shows Orco-dependent circadian modulation. (A) Example tip recordings of the spontaneous activity of one long trichoid sensillum during a period of high (top, average frequency ∼ 3 Hz) and low (bottom, average frequency ∼ 0.2 Hz) activity. **(Bi)** Spontaneous spike frequency of one long-term ORN recording during 17:7 light-dark cycles (same experiment as in **A,** colored arrows indicate respective Zeitgeber times (ZT)), indicated by the black-white bar at the top, increased during each activity phase and was low during the resting phases. Spike frequencies were averaged for each 1 h bin. **(Bii)** Mean spontaneous ORN spike frequency across all LD animals in 1 h bins. Data from the individual in **Bi** is highlighted in red. Recordings had different lengths, therefore the number of recordings used for the element-wise mean for each 1 h bin decreased with the time since the start of the recording, indicated in the top panel. **(Biii)** Mean spontaneous ORN spike frequency across all LD animals in 1 h bins during the first 48 hours revealed circadian activity. **(Ci)** The spontaneous spike frequency of one long-term ORN recording in constant darkness (DD, the expected times of lights on are represented by grey bars above) exhibited a circadian pattern. The peak activity shifted in DD due to the animal’s endogenous, free-running circadian period τ = 21.27 h. **(Cii)** Mean spontaneous ORN spike frequency across all DD animals in 1 h bins. Data from the individual in Ci is highlighted in red. **(Ciii)** Mean spontaneous ORN spike frequency across all DD animals in 1 h bins during the first 48 hours after aligning the time to the first maximum in spontaneous activity (see Methods) revealed circadian activity. **(Di)** The spontaneous spike frequency of one long-term ORN recording in constant darkness with infusion of the Orco antagonist OLC15 (orange frame) dissipated the circadian rhythm of spontaneous activity. **(Dii)** Mean spontaneous ORN spike frequency across all OLC15 animals in 1 h bins. Data from the individual in **Di** is highlighted in red. In contrast to LD and DD conditions, the spike frequency decreases over time with prolonged exposure to OLC15. **(Diii)** Mean spontaneous ORN spike frequency across all OLC15 animals in 1 h bins (mean ± SD) during the first 48 hours after aligning the time to the first maximum in spontaneous activity (see Methods). In contrast to LD and DD conditions, the circadian change in spontaneous spiking frequency disappeared. **(E)** The slopes of the linear fits to the binned spiking activity of each individual animal in the three different conditions (see Methods). Each dot indicates the slope for one experiment. The line indicates the mean. The slope for OLC15 is significantly different from control LD.

Since ORN firing patterns had a clear daily rhythm in LD both in individual animals (Figure 3Bi) and averaged across animals (Figure 3Bii, Biii), we examined whether these rhythms depended on cycling light cues or whether they persisted in constant darkness (DD), as expected for an endogenous, circadian clock-driven rhythm. In DD, the spontaneous ORN firing pattern remained rhythmic (RAIN: 8 of 12 animals), albeit with less robustness: The single peak of high spiking activity that was observed at the same ZT across several days in LD divided into multiple bouts of high spiking activity in the activity phase in 8 of 12 animals (Figure 3Ci). Also, in DD conditions, the high spiking activity phase shifted into the subjective day due to the individual endogenous circadian periods of about 23.5 ± 2.8 h (21.27 h in Figure 3Ci). Using pheromone exposure as additional Zeitgeber also did not synchronize the animals (Figure 3 – Supplement). Thus, averaging the instantaneous spiking activity across all DD animals did not depict clear circadian rhythms (Figure 3Cii). Therefore, we phase-aligned the mean frequency of spontaneous activity (see Methods). After alignment, circadian rhythms were visible across DD animals (Figure 3Ciii), comparable to the daily rhythms in LD (cf. Figure 3 Biii and Ciii).

Next, we examined whether Orco, a leaky, non-specific cation channel, is the dominant depolarizing pacemaker current of ORNs that drives circadian rhythms in spontaneous spiking activity. Infusion of the Orco antagonist OLC15 into the sensillum lymph obliterated circadian rhythms and attenuated the spontaneous activity in several, but not all experiments (RAIN: 2 of 12 animals expressed circadian rhythms) (Figure 3Di). This attenuation resulted in a linear decrease in spiking activity over several days (Figure 3Dii, Diii). Even after phase-aligning the individual experiments, no circadian rhythm was present in the average across all OLC15-treated animals (Figure 3Diii).

Because OLC15 is not membrane-permeable on its own it was infused with low concentrations (0.05%) of DMSO through the recording electrode into the sensillum lymph and, therefore, effectiveness increased over time. Hence, we fit the spiking activity of each animal and in each condition with linear regression lines over the whole recording time to compare the slopes of the fits (Figure 3E). In LD, the mean ± SD slope was 0.004 ± 0.006 Hz/h, indicating that the average spiking activity remained similar throughout the duration of the recording. In DD, the slope was −0.003 ± 0.018 Hz/h and was not different from LD. In contrast, the mean slope during OLC15 treatment was −0.007 ± 0.022 Hz/h, significantly smaller than in LD (ANOVA on ranks with Dunn’s post-hoc test; H(2) = 7.681, p = 0.021). The negative slope corroborated the finding that spiking activity decreased over time in the presence of the Orco blocker.

Activity rhythms between hawkmoths desynchronize and have only weak phase coupling in the absence of pheromone stimulation, even in LD cycles (Ghosh et al., 2024). Therefore, we first determined the percentage of animals that expressed circadian rhythmicity (period of 24 ± 4 h) for each attribute with RAIN (Figure 4A). In LD, all attributes except #AP per burst and intra-burst frequency showed daily modulation in most animals. Circadian rhythms were still present in more than half of the animals in DD for mean spiking frequency, mean burst frequency, and mean event frequency. In contrast, the attributes of only a few animals remained rhythmic with the Orco channel blocker OLC15.

**Figure 4:**
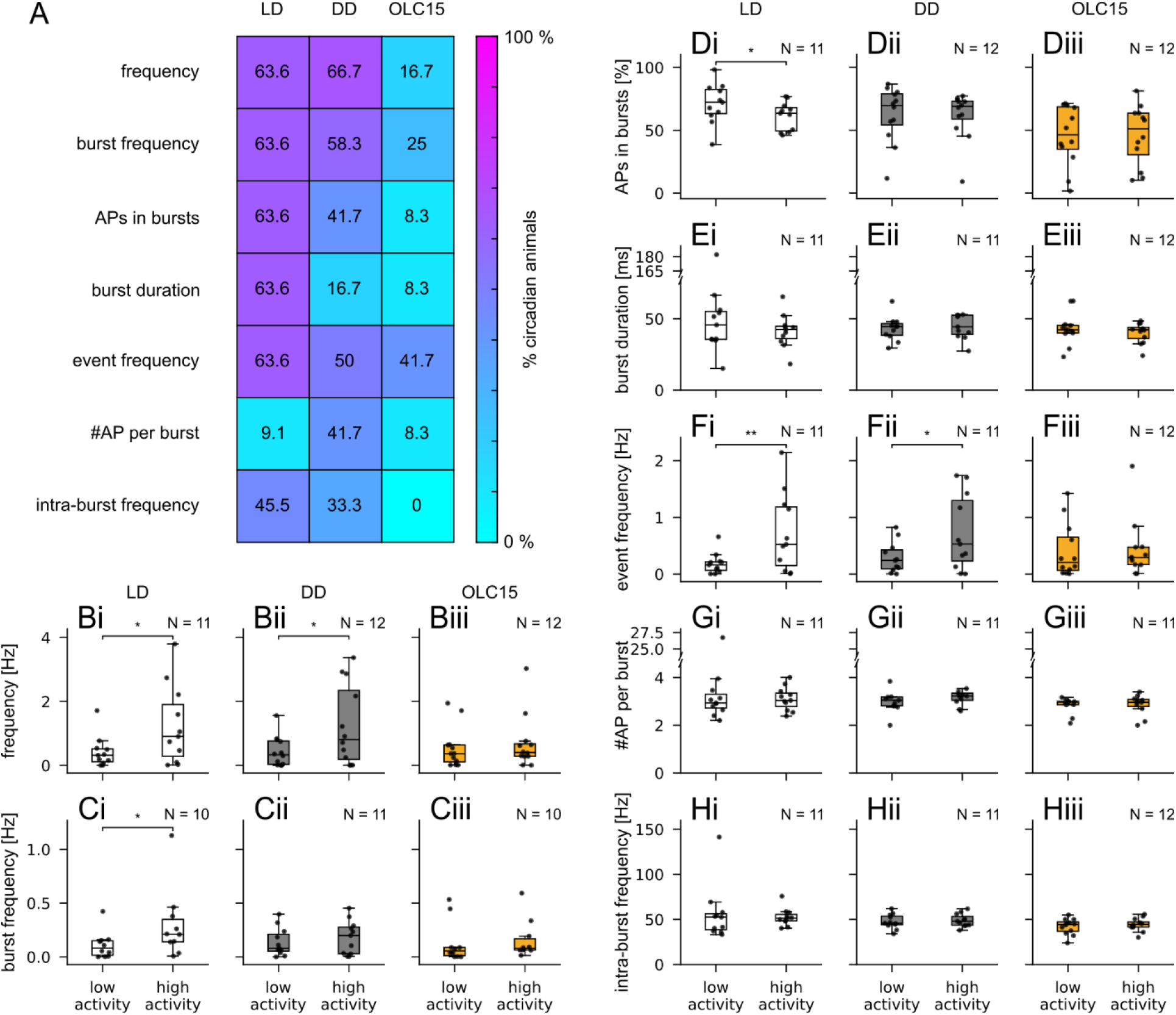
Blocking Orco removed circadian modulation of spontaneous ORN spiking patterns. RAIN analyses demonstrated significant circadian rhythmicity for most animals in most attributes of the spontaneous spiking activity in LD and DD conditions but not in OLC15 (**A**). Values and background colors indicate the percentage of animals that expressed significant circadian rhythmicity. Circadian differences of attributes were further quantified between the time windows of low vs. high activity (Wilcoxon signed-rank test, α = 0.05; **B-H**). Significant differences occurred mostly in LD (white boxes) and DD (gray boxes) conditions, but never when Orco was blocked with OLC15 (orange boxes). In both LD and DD, the mean spiking frequency was increased during high activity (**Bi, Bii**). Only in LD but not in DD, the mean burst frequency (**Ci, Cii**) increased, whereas the relative number of spikes belonging to a burst (**Di, Dii**) decreased significantly during the high activity period. Both in LD and DD the mean event frequency (**Fi,Fii**) increased significantly. Mean burst duration (**Ei, Eii**), mean number of spikes per bust (**Gi,Gii**), and mean intra-burst intervals (**Hi, Hii**) did not differ between low vs. high spiking activity in LD and DD. When blocking Orco, the attributes for low vs. high activity did not differ for any of the attributes tested (**Biii-Hiii**). In DD and OLC15conditions, activity phases were aligned as described in the Methods; thus, low activity in DD was at subjective ZT 19 and in OLC15 at subjective ZT 13, high activity was in both cases at subjective ZT 24. ZTs for low and high activity in LD were 10 and 0, respectively.

To further quantify the circadian differences, especially of the desynchronized animals, we compared different attributes of spontaneous activity between periods of high and low activity for DD conditions after phase alignment (see Methods) instead of comparing fixed ZT or circadian time (CT) intervals. Despite the high variability in ORN spiking between individuals, the mean frequency of spontaneous ORN activity was significantly higher during the high activity phase than during low activity in both LD and DD recordings (Figure 4Bi, Bii). In LD, but not in DD, the frequency of bursting increased significantly during the high activity phase (Figure 4Ci, Cii). The percentage of spikes that are part of bursts was less variable and lower during the high activity phase in LD but not significantly different in DD (Figure 4Di, Dii). The mean burst duration was less variable during the high than the low activity phase in LD, but not significantly shorter in any condition (Figure 4Ei, Eii). Furthermore, both in LD and DD, the mean inter-event frequencies were significantly higher during the high activity phase (Figure 4Fi, Fii), indicating a significantly higher overall spontaneous spiking activity during the hawkmoth’s activity phase. The mean number of spikes per burst, and intra-burst frequencies did not differ significantly between high or low activity periods in both LD and DD conditions (Figure 4Gi, Gii, Hi, Hii). The addition of Orco antagonist OLC15 deleted any significant differences of spontaneous activity found in LD and DD (Figure 4Biii-Hiii). In conclusion, the mean frequency of spontaneous spiking and the frequency of bursting expressed circadian modulation, and are both most likely controlled via a circadian clock that involves Orco.

### 5.2 Orco imposes circadian modulation on the ultradian rhythms in spontaneous ORN activity

After establishing that Orco influences the circadian changes in ORN spiking activity between the animal’s resting and activity phase, we examined if Orco also influences ORN spiking rhythms on the largely different ultradian time scales in the frequency range between 0.01-1000 Hz. In all conditions, two broad bands of ultradian instantaneous frequencies were evident: one upper band of >10 to ∼100 Hz (beta/gamma range) and a lower band between 0.1 to ≤10 Hz (delta/theta/alpha range) (Figure 5). The upper band primarily represents the frequencies of spikes within a burst, while the lower band represents single spikes and frequencies between bursts.

**Figure 5:**
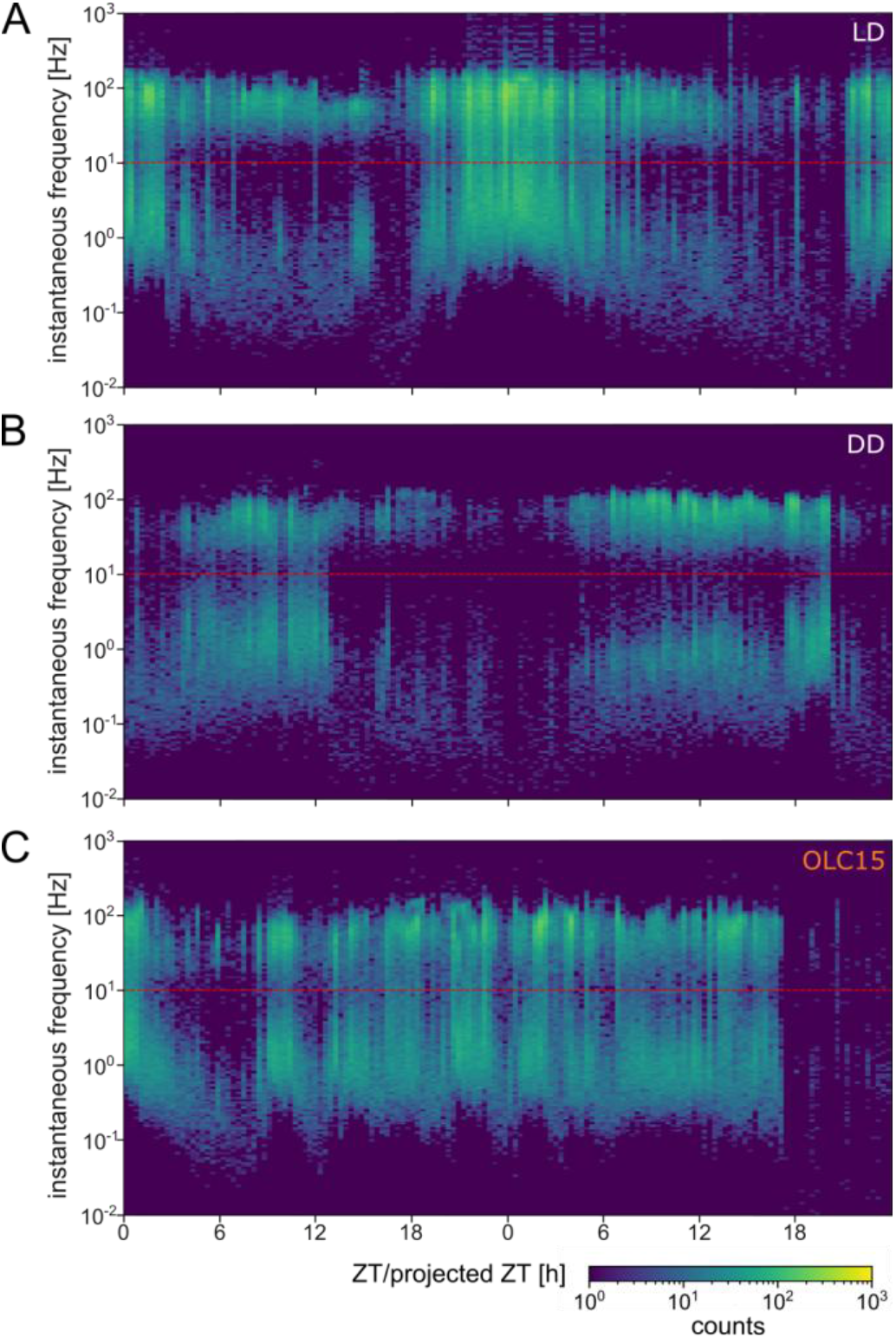
Blocking Orco removed circadian regulation of ultradian rhythms in the spontaneous ORN spiking pattern. Heat map of the instantaneous frequencies (1/ISI) over two consecutive days of long-term tip recordings of one pheromone-sensitive long trichoid sensillum under different conditions (LD **(A)**, DD **(B)**, OLC15 **(C)**) in each panel. Representative recordings from one animal in each panel. Pixel color indicates the counts of instantaneous frequencies in that respective bin. The points fall mostly in a band of high frequencies (>10 – ∼100 Hz) and low frequencies (0.01 – ∼10 Hz), with the high-frequency band representing the instantaneous frequencies of spikes within a burst and the low-frequency band the instantaneous frequency between bursts. **(A, B)** The high frequency band indicates daily and circadian modulation of frequency prevalence. In addition, the low frequency band (<0.01 Hz to ∼ 10 Hz) also displayed circadian modulation of the frequency composition. **(C)** Infusion of OLC15 deleted the circadian modulation in both frequency bands but not the ultradian rhythm of frequency prevalence.

In LD, two types of daily/circadian modulation of the ultradian frequencies occurred (Figure 5A). The upper frequency band was mostly modulated in prevalence (its frequency of occurrence) but not in its frequency range. The lower band was modulated both in prevalence and frequency range, which resulted in a sinusoidal change in the mean of the frequency band over 24 h. During the hawkmoth’s activity phase, higher frequencies of up to ∼10 Hz occurred, while frequencies dropped below 1 Hz during the resting phase. Because of this circadian increase in the lower frequency band, it merged with the high frequency band when both bands had maximum prevalence during the activity phase around ZT 0.

The circadian modulation of both frequency bands was maintained in constant darkness (Figure 5B) but phase-shifted with respect to the projected ZT due to the endogenous circadian period of each individual. Again, this indicates that a clock is modulating the spontaneous ORN spiking activity over the course of a day. The addition of the Orco antagonist OLC15 deleted any circadian modulation of either frequency band. Interestingly, the instantaneous frequencies in both bands still showed correlated changes in prevalence and merging, but now at ultradian periods only.

Additional Fourier analysis (see Methods) of spontaneous spiking activity over the course of several days revealed faster ultradian and slower infradian rhythms, ranging from 0.6 h to 38 h, in addition to the circadian (24 ± 4 h) rhythm (Figure 6Ai, Bi, Ci). Some rhythms (factorials of 24 h, like 12 h, 6 h, etc.) appeared to be harmonics of the circadian rhythm, which were not excluded in our analysis. Most animals expressed ultradian rhythms of 2-4 h in LD as well as DD conditions. Fourier confirmed the decrease of animals with circadian spiking activity in OLC15 (58.3% in DD, 33.3% in OLC15). Please note that discrepancies in the number of animals with circadian rhythms between RAIN and Fourier/wavelet analyses are due to the fundamentally different sensitivities in methodology. In brief, Fourier analysis assumes a stationary sinusoidal component, whereas RAIN as a rank-based umbrella test detects consistent raise-fall patterns, including asymmetric rhythms. Nonetheless, both methods confirmed a dramatic decrease in animals with circadian spiking patterns in OLC15. In addition, 33.3% of animals expressed infradian spiking patterns (up from 0% in DD) with OLC15. Ultradian periods could be detected in all animals in all conditions, but their distribution did not differ between conditions (1-way ANOVA; F(2, 117) = 0.87, p = 0.421). These results were corroborated by continuous wavelet analysis, which provided time-resolved spectral power across different rhythmic periods (Figure 6Aii, Bii, Cii). In control LD and DD conditions, wavelet power for ultradian periods increased during times of elevated firing rate and decreased during periods of low activity (Figure 6Aii and Bii, middle and bottom panels). This dynamic pattern indicates that ultradian rhythms were not constant over time but were instead modulated with a circadian rhythm, becoming more prominent during circadian peaks in neuronal activity. Such circadian gating of ultradian power was clearly visible in the wavelet spectrograms as alternating bands of higher and lower ultradian power aligned with daily activity cycles. Following Orco blockade with OLC15, this modulation was disrupted (Figure 6Cii middle panel): although ultradian components persisted (Figure 6Cii bottom panel), they no longer showed systematic variation with the circadian cycle, suggesting a decoupling of ultradian rhythms from circadian control.

**Figure 6:**
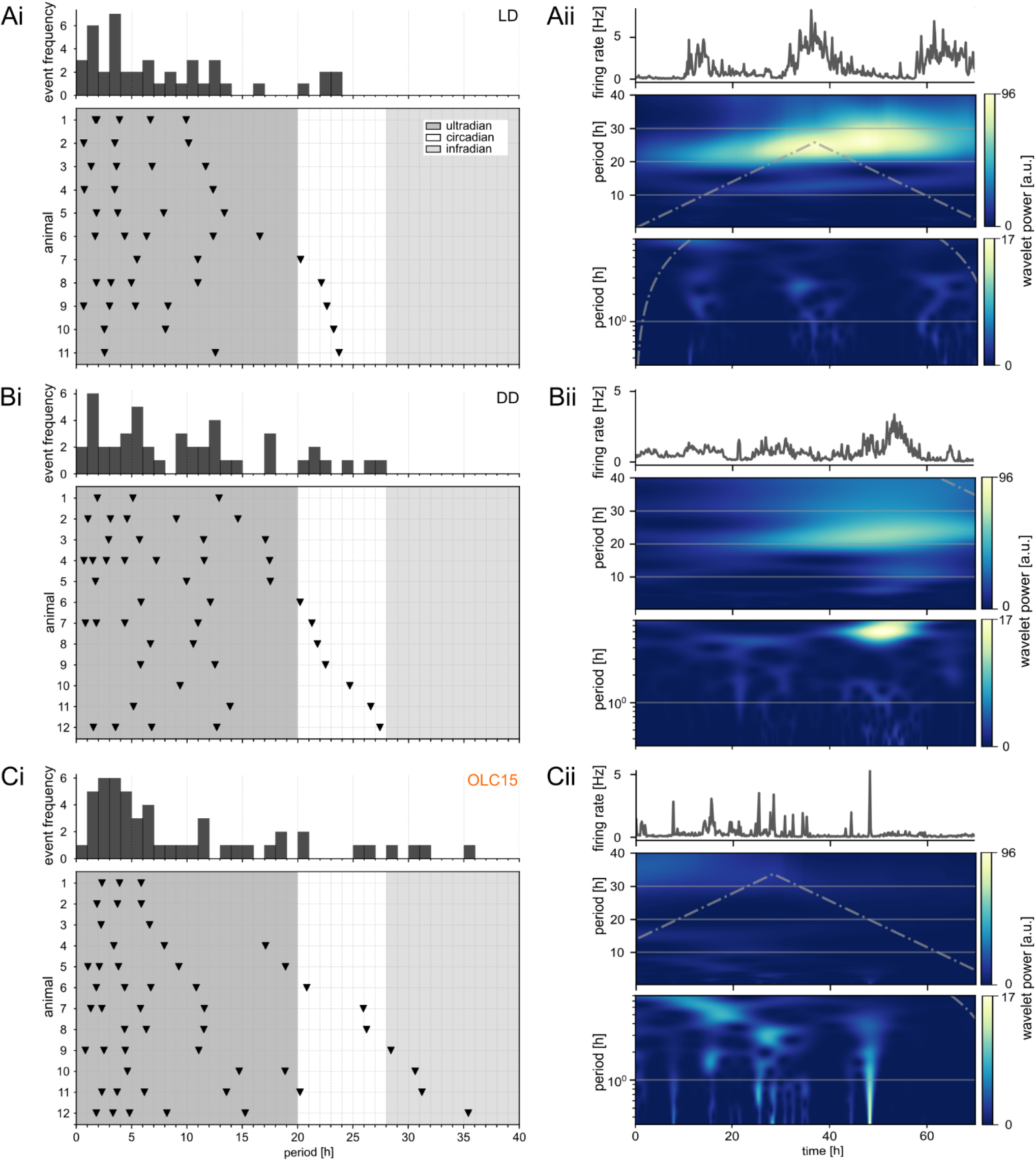
Blocking Orco affected ultradian and infradian frequencies in the spontaneous ORN firing patterns. The Fourier analysis of spontaneous spiking activity revealed rhythms with ultradian (<20 h; dark grey), circadian (20-28 h; white), and infradian periods (>28 h; light grey) in LD (**Ai**), DD (**Bi**), and OLC15 (**Ci**). Each line represents one animal where each triangular marker is at the local maximum in the frequency spectra obtained for that animal. The histograms above illustrate how often specific periods of spiking activity rhythms occurred averaged over all animals. Circadian rhythms were detected in LD (N = 5 of 11), DD (N = 7 of 10), and OLC15 (N = 4 of 12). Wavelet analysis of LD (**Aii**), DD (**Bii**), and OLC15 (**Cii**) confirmed the occurrence of multiscale periods. Example plots for one animal each, the same animals as in Figure 5. For each panel, the top plot depicts the mean firing frequency, the middle plot the wavelet power in the same period range as panels **i**, and the bottom plot the wavelet power for infradian periods up to 7 h on a log scale to highlight infradian time scales. Gray dashed lines indicate cone of uncertainty.

### 5.3 Orco expression is not under the control of the molecular TTFL clockwork

So far, our results indicated a prominent role of Orco in the circadian modulation of ORN spiking activity. Since it is the general opinion that the master circadian clock is the TTFL clock that dominates all physiological and behavioral circadian rhythms via transcriptional control, we examined whether Orco transcript abundance shows a daily rhythm, as previously found for clock genes in hawkmoth antennae (Schneider et al., 2025; Schuckel et al., 2007). However, in contrast to the circadian clock gene *tim*, qPCR of whole male hawkmoth antennae across different ZTs revealed no significant daily rhythm of Orco transcript abundance (Figure 7; ANOVA on ranks, H(5) = 6.91, p = 0.227). Thus, we suggest that Orco function is controlled via a post-translational circadian clockwork mechanism that is associated directly with the plasma membrane (Stengl and Schneider, 2024; Stengl and Schröder, 2021).

**Figure 7:**
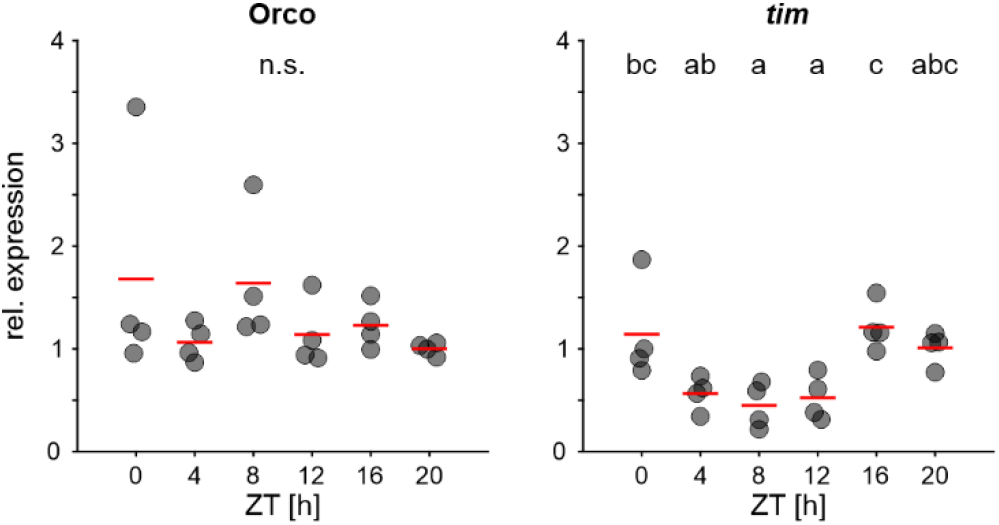
Orco is not under control of the TTFL-based molecular circadian clock. Relative expression levels of *Orco* and *tim* mRNA from male hawkmoth antennae raised under LD conditions. *Orco* expression levels did not differ significantly between ZTs (ANOVA on ranks, H(5) = 6.91, p = 0.227: N = 4 per ZT). *tim* expression levels changed significantly throughout the day (1-way ANOVA, F(5, 18) = 6.215, p = 0.002) Red lines indicate mean, *tim* served as positive control.

### 5.4 Circadian changes in Orco conductance are sufficient to modulate spiking activity in a computational model of a hawkmoth ORN

The maximum conductance of *Drosophila* Orco as a leaky ion channel is modulated by cAMP (Getahun et al., 2013; Martín et al., 2001; Sargsyan et al., 2011; Wicher et al., 2008). In hawkmoth antennae, cAMP levels oscillate throughout the day and are involved in sensitizing the ORNs to pheromone (Dolzer et al., 2021; Flecke et al., 2010; Flecke and Stengl, 2009; Schendzielorz et al., 2014, 2015). Hence, cAMP modulation of Orco conductance is a putative mechanism for the Orco-dependent circadian modulation of ORN spiking activity, integrating Orco in a circadian PTFL-based membrane clockwork (Stengl and Schneider, 2024).

To test the hypothesis that Orco is a cAMP-dependent, leaky ion channel, we constructed an ORN model (Methods and Supplementary Material) and tuned the parameters to provide a quantitative match with the biological data. The developed model provides a unified description of experimental observations. For instance, the ORN model yields irregular spiking activity with a bimodal distribution, consisting of both bursts and isolated spikes, successfully reproducing the experimental recordings (Figure 8A). This suggests a unified mechanism underlying the initiation of both phenomena: If the pacemaker-driven depolarization reaches the Na^+^ threshold, a firing process begins, and at this point randomness determines if other slower ion channels responsible for bursting are sufficiently activated to sustain a burst of several spikes.

**Figure 8:**
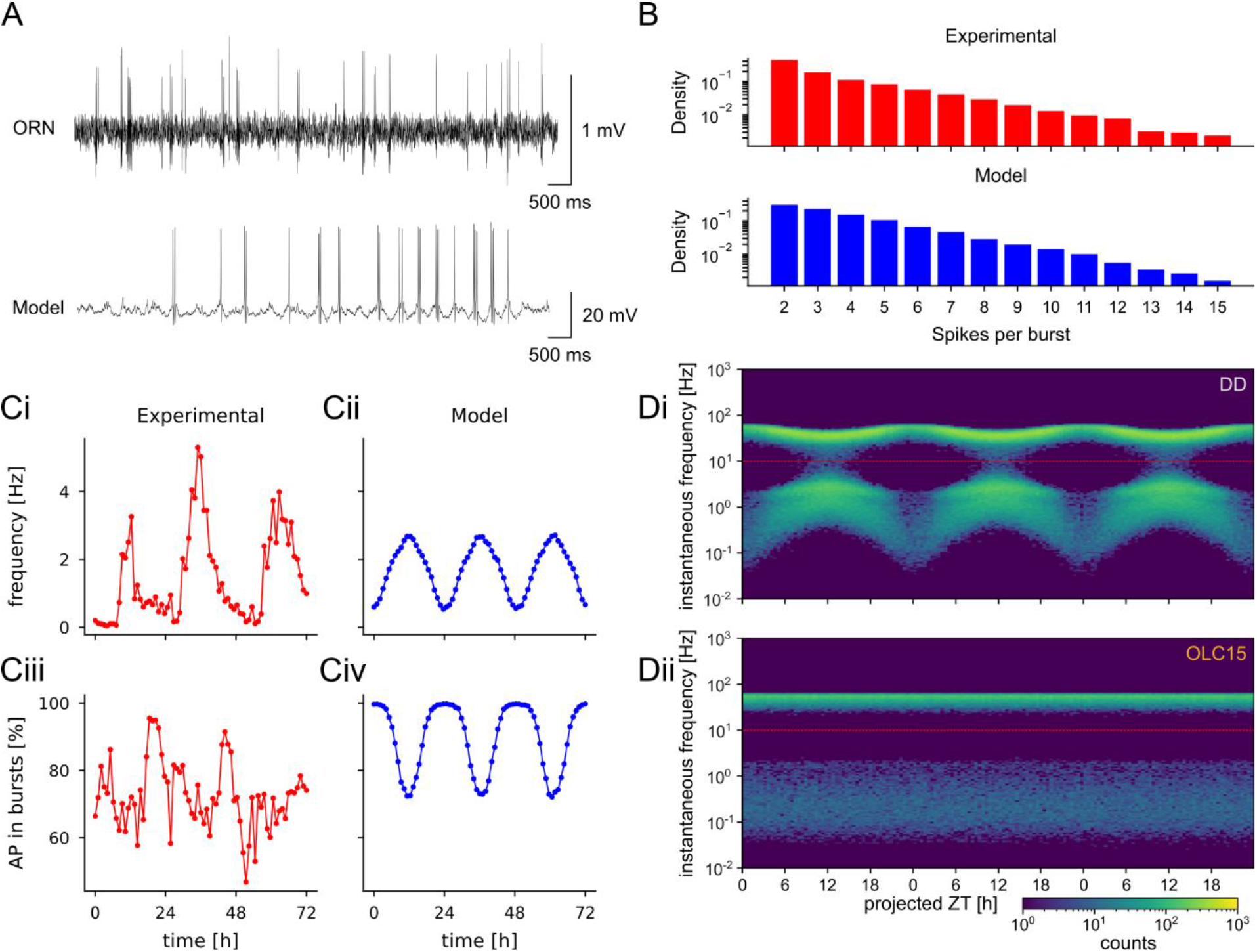
The model adequately reproduces the firing pattern of the biological ORN. **(A)** The model activity (bottom trace) displays both isolated spikes and bursts of variable length as in the original recording (top trace). The simulated trace shows the membrane potential (an intracellular variable), while the biological data were obtained with extracellular electrodes. **(B)** Semi-logarithmic plot of the distribution of spikes per burst of an experimental LD recording (top) and simulated model (bottom) of spontaneous action potential activity of pheromone-sensitive neurons. The experimental distribution corresponds to the 1-hour period of highest firing frequency in the recording; the simulated distribution was obtained from a separate 1-hour simulation at fixed high Orco conductance. Both the model and the recordings display exponential decrease of the count’s density as a function of the burst length, shown as a linear decrease in the logarithmic scale. **(C)** Comparison of experimentally recorded and model-predicted circadian modulation of spiking attributes. Top panels show mean spike frequency (**Ci**, experimental; **Cii**, model), whereas bottom panels show the percentage of spikes belonging to bursts (**Ciii**, experimental; **Civ**, model). The simulated model reproduced the circadian dynamics observed in the experimental recordings, demonstrating good agreement between model and biological data. **(D)** Heat map of the simulated model, x-axis shows time (days), y-axis shows instantaneous frequency (Hz), and pixel color indicates count density on a logarithmic scale. The instantaneous frequencies can be roughly divided in a band of high frequencies (top, ∼50 – 80 Hz) and low frequencies (bottom, ∼0.1 – 5 Hz), with the high-frequency band representing the instantaneous frequencies of spikes within a burst and the low-frequency band the instantaneous frequency between bursts. **(Di)** Simulated spikes with circadian regulation by the Orco channel; the inter-burst frequencies varied widely with circadian rhythmicity while the frequencies within burst remained approximately constant. The simulated results also reproduced the “merging” effect between the two bands when the bottom one approaches the top one. **(Dii)** Simulated spikes without circadian regulation of Orco had no circadian oscillations.

The model successfully reproduces the circadian modulation of spiking activity: the maximum firing rate occurs when the Orco conductance is maximal, whereas the maximum percentage of spikes that are part of bursts correlates with the lowest Orco conductance, resulting in antiphase oscillations, as were observed in the experimental recordings (Figure 8C).

The frequency prevalences in the model over several days clustered in two distinct frequency bands: an upper band that corresponds to the intra-burst frequencies and a lower band, corresponding to inter-burst frequencies (Figure 8D), in agreement with the biological experiments (Figure 5A, B). When circadian oscillations in Orco conductance were included, inter-burst frequencies showed pronounced circadian variations, whereas intra-burst frequencies remained relatively constant, reminiscent of the biological data. Notably, the simulation captured the experimentally observed “merging” effect, where the lower frequency band approaches the upper band during high-activity periods (Figure 8D).

This merging of spiking and bursting frequencies suggests that the system approached a bifurcation condition coming from the distribution of spikes per burst (Figure 8B), computed from the 1 h period of highest firing frequency in an experimental recording and from a separate 1 h simulation at fixed high Orco conductance. In both cases, the distribution followed a decreasing exponential trend – in contrast to the constant value or bounded range expected from a standard bursting neuron model – consistent with a system operating near a bifurcation point. The observation that the circadian modulation of spiking activity requires only one parameter, namely changes in the Orco conductance, to describe the circadian effects on the ORN spiking pattern indicates the crucial role of the Orco channel in circadian control.

### 5.5 cAMP regulates the circadian function of Orco

Our computational model predicted that the daily/circadian oscillations in cAMP levels (Schendzielorz et al., 2015) control the circadian modulation of spontaneous spiking activity of hawkmoth ORNs. To test this prediction in hawkmoth ORNs, we increased cAMP levels by infusing 8-Br-cAMP-AM through the recording electrode into the sensillar lymph (Figure 9) at ZT 1-3, when naturally occurring cAMP levels are lowest (Schendzielorz et al., 2015) to maximize the effect. This significantly increased the mean frequency of the spontaneous spiking activity (Figure 9A; Wilcoxon signed-rank test Z = −2.701; p = 0.004). To check whether this increase in mean frequency was caused by direct or indirect interaction of cAMP with the Orco channel, we infused the Orco antagonist OLC15 together with 8-Br-cAMP-AM through the same recording electrode. Blocking Orco prevented a significant change in the spontaneous spiking frequency via the cAMP analogue (Figure 9B; paired t-test t(9) = −1.446; p = 0.182). This further corroborates cAMP-dependent modulation of the circadian pacemaker channel Orco.

**Figure 9:**
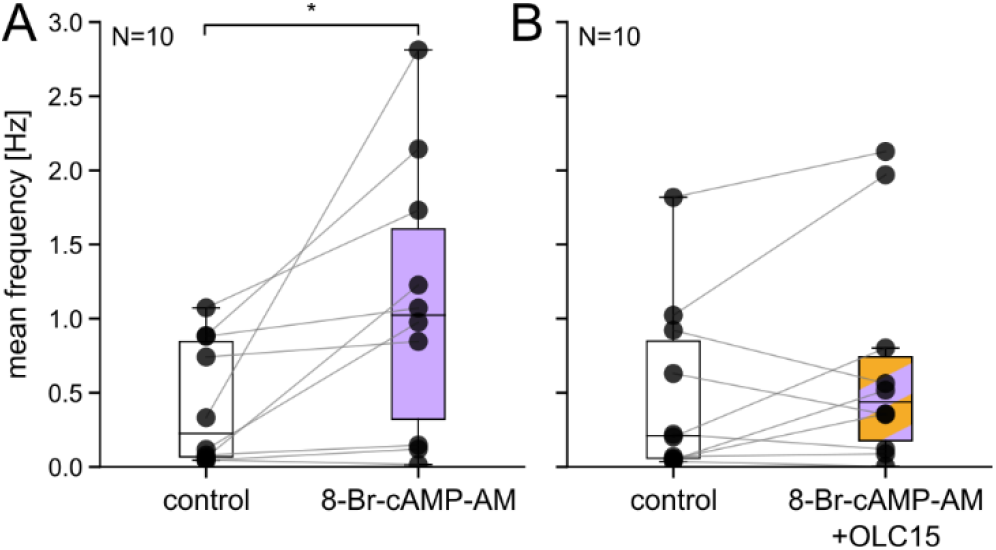
cAMP modulates open-time probability of Orco in M. sexta. Perfusion of 8-Br-cAMP-AM significantly increased the spontaneous spiking activity at the end of the activity phase (ZT 1-3; **A**; Wilcoxon signed-rank test Z = −2.701; p = 0.004). However, activity did not increase when 8-Br-cAMP-AM was added together with the Orco blocker OLC15 (**B**; paired t-test t(9) = −1.446; p = 0.182). Electrodes were filled with sensillum lymph ringer containing 0.1% and 0.15% DMSO as control in **A** and **B**, respectively. Data from individual animals are connected by lines.

## 6 Discussion

Searching for daily and circadian rhythms of Orco-dependent membrane potential control in olfactory receptor neurons (ORNs) of *M. sexta*, *in vivo* long-term tip recordings of pheromone-sensitive long trichoid sensilla of male hawkmoth antennae were performed. We detected interlinked circadian and ultradian frequencies in the spontaneous spiking activity, indicative of underlying membrane potential oscillations in ORNs. We found that the olfactory receptor co-receptor (Orco) acts as a pacemaker channel that controls the daily/circadian modulation of the spiking activity without affecting the ultradian gamma frequency range that is relevant for the primary fast, phasic pheromone response (Dolzer et al., 2003; Nolte et al., 2016; Schneider et al., 2025). Furthermore, since OLC15-dependent block of Orco did not delete ultradian action potential activity, other additional pacemaker channels must be present in ORNs that possibly control different frequency ranges of ultradian membrane potential oscillations. Since qPCR experiments at different ZTs showed that Orco transcription was not directly controlled by the circadian TTFL clock present in hawkmoth ORNs, we concluded that the daily/circadian control of Orco function is obtained at the post-translational level. Our computational model predicted that cAMP-dependent post-translational modulation of Orco conductance is sufficient to account for our electrophysiological findings if cAMP levels vary in a daily/circadian rhythm. Daily rhythms in cAMP levels are confirmed in hawkmoth antennae (Schendzielorz et al., 2015). Furthermore, here we provided evidence that cAMP analogs Orco-dependently control spontaneous activity of hawkmoth ORNs. Thus, we suggest that, colocalized in a membrane signalosome in the ORN soma compartment, constitutively depolarizing Orco homomeric ion channels generate ultradian membrane potential oscillations via delayed activation of hyperpolarizing K^+^ channels (Figure 10). The ultradian membrane potential oscillations express a daily/circadian rhythm with higher frequencies during the hawkmoth’s activity phase at night compared to the rest phase during the day. Thus, as novel hypothesis we suggest that the soma signalosome constitutes an endogenous, plastic, post-translational feedback loop (PTFL) clockwork, controlling membrane potential oscillations with Orco as 2^nd^ messenger-dependent control hub that is coupled to, but not commanded by, the circadian TTFL clock in the nucleus.

**Figure 10:**
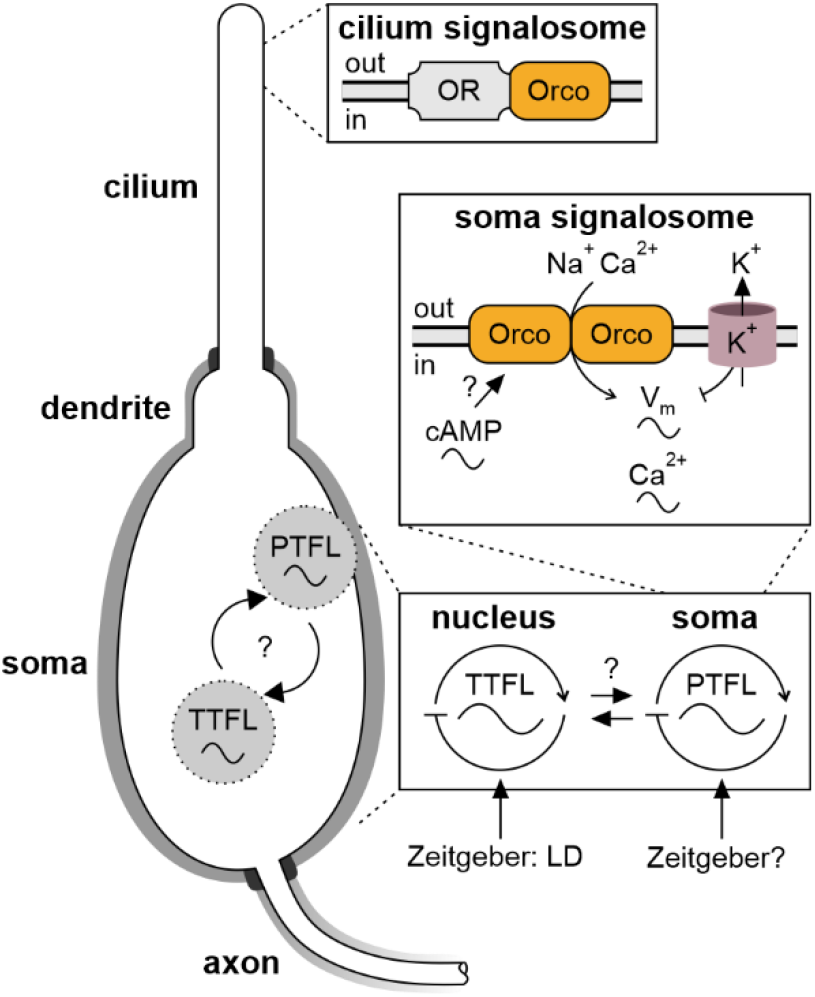
A signalosome assembling an endogenous multiscale PTFL membrane clock in the soma of hawkmoth olfactory receptor neurons (ORNs) with 2^nd^ messenger-dependent Orco as central control hub is proposed to mediate membrane potential rhythms. Schematic of an antennal pheromone-sensitive ORN that is strictly compartmentalized by tight junctions (thick black borders) into three functionally separated membrane compartments: cilium, soma/dendrite, and axon. Soma/dendrite and axon are wrapped by supporting cells (grey background), ensuring common extracellular ionic concentrations for both membrane compartments. Thus, only the cilium is exposed to the sensillum lymph that contains high (∼200 mM) K^+^ concentration. The ciliary membrane comprises a specialized membrane signalosome devoted to highly sensitive pheromone detection. In contrast, a signalosome in the soma/dendrite controls the membrane potential and intracellular Ca^2+^ concentration. Sustained depolarization via opening of cation-permeable, homomeric Orco pacemaker channels in the soma are counteracted by delayed voltage-gated hyperpolarizing K^+^-channels, together generating ultradian oscillations in the membrane potential with interlinked oscillations in intracellular Ca^2+^. Circadian/daily rhythms in cAMP concentrations mediating Orco conductance, next to Ca^2+^, superimpose a circadian rhythm on prevalence of ultradian oscillations. Orco is not under direct circadian transcriptional control by the circadian TTFL clock in the nucleus that is entrained to the LD cycle, thus being part of a PTFL membrane clock entrained to other Zeitgebers. PTFL- and TTFL-clocks are coupled by not-yet identified mechanisms. LD: light dark cycle; OR: odor receptor, Orco: olfactory receptor coreceptor, PTFL: post-translational feedback loop clock; TTFL: transcriptional-translational feedback loop clock; V_m_: membrane potential; sine waves: oscillations.

The expression of a 2^nd^ messenger-dependent leaky pacemaker channel in ORNs, which does not require daily transcription, would constitute an economic and rapidly adjustable mechanism of active sensing. As part of a PTFL membrane clock, Orco could orchestrate endogenous membrane potential oscillations that are tuned to detect and resonate with behaviorally relevant temporal patterns of Zeitgeber signals, such as the circadian and ultradian pheromone fluctuations in the insect’s environment that are detected in the cilia of ORNs (Stengl and Schneider, 2024; Stengl and Schröder, 2021). The present study provides first evidence for a systems view of biological timing that involves multiscale PTFL membrane clocks linked to the TTFL clock in insect sensory neurons, instead of the prevailing hierarchical view that assumes all endogenous circadian oscillations are outputs of the genetic circadian TTFL clock (Stengl and Schneider, 2024; Stengl and Schröder, 2021). Future studies will challenge our novel hypothesis.

### 6.1 Orco in the ORN soma compartment acts as a pacemaker channel that controls the membrane potential oscillations underlying spontaneous spiking activity

The resting membrane potential of most silent neurons lies close to the negative equilibrium potential for K^+^ ions, mainly due to potassium leak channels and the driving force by the 10x higher intracellular K^+^ concentration (Goldstein et al., 2001; Hille, 2001; Patel and Honoré, 2001; Talley et al., 2001). Thus, spontaneously active neurons require the expression of Na^+^ and/or Ca^2+^ permeable leak or pacemaker channels that open at the hyperpolarized resting membrane potential, thus depolarizing the neuron via influx of cations along their concentration gradients (Bose et al., 2014; Cochet-Bissuel et al., 2014; Das et al., 2016; Golowasch et al., 2017; Lüthi and McCormick, 1998; Ratliff et al., 2021; Robinson and Siegelbaum, 2003; Sharma et al., 2023). When the pacemaker-driven depolarization reaches spike threshold, voltage-gated Na^+^ and K^+^ channels in the axon promote spiking. Furthermore, voltage-dependent Ca^2+^ channels allow for depolarization-dependent Ca^2+^ influx. Voltage- and Ca^2+^-gated hyperpolarizing K^+^ channels then repolarize the neuron to negative membrane potentials, re-starting the cycle. Furthermore, different fine-tuned mechanisms of homeostasis re-establish intracellular low (pico- to nanomolar range) Ca^2+^ concentrations, preventing necrosis. Thus, membrane potential oscillations with interlinked oscillations in intracellular Ca^2+^ levels drive spontaneous spiking activity, which depends on the type of pacemaker channel and antagonistic ion channels present.

In insect ORNs, at least two types of pacemaker channels exist: the highly conserved, slow, leaky Orco, which we focus on in this study, and an inversely voltage-dependent HCN type channel (Butterwick et al., 2018; Dolzer et al., 2021; Kodirov, 2022; Lee and MacKinnon, 2017; Mandala et al., 2025; Nolte et al., 2016, 2013; Sato et al., 2008; Stengl and Funk, 2013; Stengl and Schneider, 2024; Stengl and Schröder, 2021; Wicher et al., 2008; Wicher and Miazzi, 2021). When expressed *in vitro* in heterologous expression systems, *Drosophila* as well as *Manduca* Orco homo- and heteromers assemble into leaky, non-specific cation channels with a reversal potential around 0 mV, being constitutively permeable for Na^+^, K^+^, and Ca^2+^ (Nolte et al., 2016, 2013; Sato et al., 2008; Stengl and Funk, 2013; Wicher et al., 2008; Wicher and Miazzi, 2021). Thus, while this study did not aim to unravel the overall generation of spontaneous activity in ORNs, here, we examined whether the leak channel Orco in hawkmoth ORNs is the core pacemaker channel responsible for membrane potential oscillations that drive spontaneous circadian activity (Stengl, 2010; Stengl and Schneider, 2024).

Orco homo- or heteromers are likely to have different localizations and functions in the distinct, sealed-off membrane compartments of ORNs that face different ionic environments: the cilium (also termed “outer dendrite”), and the soma / (“inner”) dendrite (Stengl and Funk, 2013; Stengl and Schröder, 2021). We propose for *Manduca*, that homomeric Orco pacemaker channels, as core element of a PTFL membrane clock, are restricted to the soma/dendrite compartment of ORNs, where they face the same extracellular cation concentrations as other brain neurons, with low K^+^, high Na^+^, and high Ca^2+^ concentrations (Stengl and Schneider, 2024; Stengl and Schröder, 2021). In contrast, we expect *Manduca* Orco heteromers to serve as obligatory chaperons to localize ORs to cilia and maintain them in the ciliary membranes, as proven in *Drosophila* (Benton et al., 2006). Since the ciliary compartment faces very high concentrations of K^+^ in the sensillum lymph, matching intracellular K^+^ levels (Kaissling and Thorson, 1980; Steinbrecht, 1992), we propose that Orco channels in the ciliary membrane remain closed. If Orco homo- or heteromers in cilia would constitute leaky pacemaker channels, their K^+^ permeability would clamp the ciliary membranes at 0 mV, due to the high extracellular K^+^ concentration, strongly reducing the driving force of pheromone-dependent receptor currents, severely compromising pheromone sensitivity (Stengl and Funk, 2013). Accordingly, it was shown previously in hawkmoth ORNs that Ca^2+^ channels, most likely of the TRP ion channel family, serve as transduction channels for the primary events of the highly sensitive pheromone transduction, and not a non-specific cation channel like Orco (Gawalek and Stengl, 2018; Nolte et al., 2016, 2013; Stengl, 1994, 1993).

In support of the specific localization of Orco homo- or heteromers, Orco antisera labeled Orco in two of the three sealed off ORN membrane compartments in both *Drosophila* (Benton et al., 2006) and *Manduca* (Nolte et al., 2016): in the membrane of the soma/dendrite compartment and in the membrane of the cilium, but not in the axons. In contrast, antisera against an OR in *Drosophila* detected ORs accumulated only in ciliary membranes but not in the other compartments of ORNs (Benton et al., 2006). Thus, the localization of OR-Orco heteromers in cilia confirm that Orco is an obligatory coreceptor and chaperone of ORs localizing and maintaining the OR-Orco complexes in the ciliary membrane (Benton et al., 2006). Since antisera against *Manduca* ORs are not available, we do not know for hawkmoths whether the pheromone-sensitive OR-Orco heteromers are located to ciliary membranes, while Orco homomers accumulate in the soma/dendrite compartment (Fandino et al., 2019). Nevertheless, Orco is likely to assemble as homomeric complexes in wild-type hawkmoth ORNs because Orco expression starts during development, coinciding with the beginning of spontaneous activity, well before ORs are present. Furthermore, the Orco agonist VUAA1 activates Orco in primary cell cultures of maturing ORNs before they become pheromone sensitive (Nolte et al., 2016; Schweitzer et al., 1976).

In summary, we propose that Orco homomers are specifically located to a signalosome in the soma/dendrite compartment of ORNs (Figure 10), controlling the ORN’s membrane potential. Because the Orco-dependent ultradian and circadian membrane potential oscillations are maintained under constant conditions, they are generated by an endogenous multiscale clock. Thus, we predict that a distinct signalosome with Orco homomers in the soma, but not in the cilia, serves as endogenous self-assembling multiscale PTFL membrane clock for the circadian control of the membrane potential. We hypothesize that this distinctly located PTFL membrane clock is coupled to, but not dominated by, the TTFL clock in the hawkmoth ORNs to allow for fast, plastic regulation of the membrane potential (Schuckel et al., 2007; Stengl and Schneider, 2024).

### 6.2 Orco is hypothesized to be a second messenger-dependent control hub of a PTFL membrane clock that regulates circadian membrane potential oscillations in hawkmoth ORNs

Focusing on the signalosome in the ORN soma compartment, our novel hypothesis predicts that Orco is a 2^nd^ messenger-dependent control hub of a somatic multiscale PTFL clock. In accordance with this hypothesis, different 2^nd^ messenger cascades regulate *Drosophila* Orco via a Ca^2+^/calmodulin binding site and via phosphorylation/de-phosphorylation at multiple sites that are conserved between species (Cao et al., 2016; Getahun et al., 2016; Guo and Smith, 2017; Mukunda et al., 2016, 2014; Sargsyan et al., 2011). In contrast to our current focus of Orco homomer function in *Manduca*, focus and data interpretation in *Drosophila* lies on the function of Orco heteromers. In *Drosophila*, Orco’s Ca^2+^/calmodulin binding site controls the localization of the OR-Orco complex in the ciliary membrane of ORNs, provides for Ca^2+^-dependent negative feedback control of Orco conductance, and depends on previous odor stimulation (Bahk and Jones, 2016; Cao et al., 2016; Guo and Smith, 2022; Jain et al., 2021). After phosphorylation of Orco by protein kinase C (PKC), *Drosophila* Orco directly or indirectly becomes cyclic nucleotide-dependent, with cGMP and cAMP increasing Orco-dependent depolarization and Ca^2+^ influx (Getahun et al., 2013; Wicher et al., 2008). However, it is not yet resolved in *Drosophila* in which of the different sensilla compartments and at what time of day Orco homo- or heteromers are controlled via 2^nd^ messengers and how this control is regulated.

Also, in *Manduca* the 2^nd^ messenger-dependent modulation of Orco is not yet understood. However, due to Orco’s highly conserved structure we assume Ca^2+^/calmodulin-, PKC-, and cyclic nucleotide-dependent control of hawkmoth Orco localization and/or open-time probability, likely similar as described for *Drosophila* Orco. It is very likely that the conserved structure of Orco with its manifold 2^nd^ messenger-modulation sites designate it as a hub for intersecting 2^nd^ messenger control in different insect species. Since blocking Orco in *M. sexta* decreased the spontaneous spiking activity in ORNs and deleted its circadian modulation, Orco is apparently the main target for circadian clock control modulating circadian membrane potential rhythms. Furthermore, the ultradian rhythms in spontaneous activity appear to be controlled by additional pacemaker channels that are not focus of the current study. Since Orco transcript abundance was constant throughout the day, Orco is not controlled directly by the circadian TTFL clock. Rather, open-time probability and/or membrane localization of Orco appear to be controlled in a circadian manner via post-translational mechanisms. Because daily rhythms in cAMP levels were found in hawkmoth antennae (Schendzielorz et al., 2015), our modeling predicted that the most parsimonious explanation for Orco’s circadian control of the membrane potential is Orco’s regulation via daily rhythms of cAMP levels. In support of this hypothesis, we demonstrated that the significant increase of spontaneous spiking activity caused by 8-Br-cAMP application is canceled with co-application of the Orco blocker OLC15. Furthermore, our modeling studies showed that this post-translational circadian control of Orco via daily/circadian levels of cAMP is sufficient to explain our physiological findings. Thus, we hypothesize that a PTFL membrane clock (Figure 10), linked to the TTFL nuclear clock, determines the circadian oscillations in pheromone sensitivity and temporal resolution of ORNs via cAMP- and Orco-dependent membrane potential control (Dolzer et al., 2021, 2008; Flecke et al., 2010, 2006; Redkozubov, 2000; Schendzielorz et al., 2015, 2012; Ziegelberger et al., 1990).

Our novel hypothesis predicts that a signalosome in the soma membrane of hawkmoth ORNs assembles a circadian PTFL membrane clock controlling daily/circadian rhythms in the membrane potential and Ca^2+^ concentration, comprising Orco as central control hub. Via unknown mechanisms that still need to be resolved, circadian oscillations in cAMP levels are generated in ORNs that are linked to Orco. As is characteristic for a clock (Stengl and Schneider, 2024), the PTFL membrane clock comprises positive feedforward elements that upregulate the ORN’s depolarization and intracellular Ca^2+^ levels, which are then sequentially downregulated via delayed negative feedback elements/mechanisms. Thus, the PTFL clock generates oscillations in the membrane potential coupled to strictly compartmentalized oscillations in intracellular Ca^2+^ levels. This predicted soma signalosome comprises the pacemaker channel Orco as a central hub that is controlled via changes in voltage, Ca^2+^, and cAMP levels that can be affected by multiple parallel pathways in the ORN. We propose a parsimonious model of the PTFL clock, generating multiscale membrane potential- and Ca^2+^ oscillations linked to cAMP oscillations. Since it is a PTFL clock, it does not require circadian degradation and circadian transcription/translation of its central constituents. Nevertheless, we predict coupling between the multiscale PTFL membrane clock and the TTFL circadian clock in the ORN’s nucleus to obtain stably synchronized circadian rhythms. As likely mechanism of coupling we predict that Ca^2+^- and cAMP-dependent kinases interlink both types of circadian clocks, thereby obtaining robust and at the same time flexible interlinked cellular rhythms (Figure 10). This system of interlinked circadian clocks are hypothesized to determine changing physiological setpoints of the ORN during circadian sleep-wake cycles (Stengl and Schneider, 2024).

### 6.3 Circadian and ultradian rhythms in spontaneous ORN activity resemble the physiologically relevant time scales for active sensing

Moths have daily rhythms in pheromone sensitivity and temporal resolution controlled via antennal circadian clocks (Flecke et al., 2010; Flecke and Stengl, 2009; Merlin et al., 2007, 2006; Schendzielorz et al., 2015; Schuckel et al., 2007; Silvegren et al., 2005). The increased pheromone sensitivity and temporal resolution during the activity phase enhances the male’s chance to detect and locate a calling female. In response to brief pheromone pulses, independent of Orco, pheromone concentration is encoded in the frequency of the first, phasic burst of spikes of the characteristic triphasic spiking sequence of ORNs (Dolzer et al., 2003; Nolte et al., 2013; Schneider et al., 2025). Because blocking Orco does not affect the ultradian gamma frequency band of spontaneous activity that correlates with the frequency ranges of the stimulus concentration-encoding the first, phasic pheromone response, the current finding further confirms that Orco is not the primary transduction channel for pheromone detection (Dolzer et al., 2003; Nolte et al., 2016, 2013). Nevertheless, Orco partakes in pheromone transduction during the later and longer-lasting time window after pheromone stimulation. Mediated through the cAMP-dependent increase in Orco conductance during the activity phase, the ORN membrane potential would oscillate pheromone-dependently at higher ultradian frequencies and with larger amplitude from a lower negative baseline resting potential. Thereby, the ORN’s pheromone sensitivity and sampling frequency of pheromone pulses could be maximized while males fly upwind in search for females.

The upregulation of neural activity during wakefulness/alertness seems to be ubiquitous among different species. For example, the synchronously recorded neurons in human EEG show fast ultradian oscillations in the alpha, beta, and gamma frequency ranges during alertness and slow ultradian oscillations in the delta and theta ranges during rest and sleep. Similarly, in the lower frequency band, the spontaneous spiking activity of hawkmoth ORNs changes from below delta during rest to theta/alpha ranges during their activity phase. While the gamma frequency band comprises spike frequency within a burst and matches the first phasic pheromone response (Dolzer et al., 2003), the spontaneous spiking activity in the theta/alpha range matches the maximal temporal resolution of pheromone pulses of up to 10 Hz (Marion-Poll and Tobin, 1992), being ideally suited to tune ORNs for active sensing. Even though oscillations in the field potential of electroantennogram responses to odor stimulation can reach 100 Hz in different insects (Szyszka et al., 2014), these frequencies most likely do not encode the temporal resolution of physiological odor responses. Apparently, they reflect the astounding adaptive ability of antennal ORNs to synchronize their endogenous membrane potential oscillations across the antenna to a broad range of stimulation frequencies. Pheromone pulse resolution of about 100 Hz was never reported in single olfactory sensilla recordings, and in behavioral experiments male moths stop their upwind mate search if pheromone pulses exceed 30 Hz, apparently due to adaptation (Baker et al., 1985; Bau et al., 2005, 2002; Tripathy et al., 2010).

In summary, Orco-dependent circadian oscillations in the ORN membrane potential are ideally suited to autonomously, fast, and flexibly tune sensitivity and temporal resolution of the pheromone-sensitive ORNs to the hawkmoth’s rest-activity cycle and to acute physiological requirements, synchronizing male and female mating behavior. Here, we have demonstrated how modulation of the leaky pacemaker channel Orco can achieve these circadian oscillations. Whether cAMP levels are under control of the TTFL clock or oscillate based on Ca^2+^ - and/or octopamine concentration oscillations, being part of a stand-alone PTFL clock, remains to be studied.

### 6.4 Stochastic ion channel dynamics and circadian modulation of Orco conductance explain modeled ORN firing patterns

Our modeling results provide mechanistic insight into the irregular spontaneous bursting activity of pheromone-sensitive ORNs in *M. sexta*, highlighting stochastic ion channel dynamics and circadian modulation of ion conductance that together shape spiking patterns. By employing a Langevin approximation of the stochastic HH formalism, we demonstrate that the intrinsic noise arising from the random gating of ion channels is sufficient to generate both isolated spikes and bursts of spikes, as observed experimentally. Our model effectively recapitulates the spike distributions found *in vivo* and accounts for the role of channel noise as a critical determinant of ORN firing patterns.

A key contribution of our model is the incorporation of the Orco ion channel as a pacemaker component, with a conductance that varies linearly with cAMP concentration. We show that post-translational circadian modulation of Orco conductance alone is sufficient to produce the observed time-of-day dependent changes in spike distribution. This finding suggests that the rhythmic expression of cAMP levels across the circadian and/or daily cycle may act as a primary driver of circadian and diurnal changes in ORN excitability, without requiring additional external inputs or modulation of other ion channels.

Importantly, our results reveal that the same set of ion channels can support the emergence of both single action potentials and complex bursting behaviors purely through the stochastic variability of their gating kinetics. This unifying explanation reduces the need for invoking separate biophysical mechanisms or circuit-level inputs to explain these distinct firing modes. For simplicity, the circadian variation in cAMP concentration was modeled using a sinusoidal function. While this approximation is sufficient for capturing the qualitative features of the rhythmic modulation, future work could enhance the biological realism of the model by coupling Orco conductance dynamics to an endogenous circadian oscillator, such as a Goodwin-type TTFL. Incorporating this layer would allow for a more detailed exploration of how endogenous clock components regulate cAMP synthesis and degradation. Additionally, further extensions could include downstream regulatory or coupling mechanisms such as phosphorylation and dephosphorylation events mediated by kinases and phosphatases under circadian control, potentially contributing to fine-tuned modulation of ion channel function over the daily cycle. Together, these findings demonstrate the value of a biophysically grounded, noise-inclusive modeling framework for understanding complex temporal patterns in sensory neuron activity and open the door for future integrative models linking different types of endogenous circadian clocks to cellular excitability.

## Acknowledgements

We thank Dr. Dieter Wicher (Max Planck Institute for Chemical Ecology, Jena, Germany) for generously providing OLC15, and Rishaban Radhakrishnan for help with Python codes. We thank Prof. Hülya Altuntaş for helpful discussions. Dr. Pablo Rojas current affiliation: Interdisciplinary Institute for Scientific Computing, Heidelberg University, Germany C Clinical Psychology, Central Institute of Mental Health, Medical Faculty Mannheim, Heidelberg University, Germany

## Author contributions

Conceptualization: AV, MF, MEG, MS; data curation: AV, MF, YC, PR, KS, ACS, MEG, MS; formal analysis: AV, MF, YC, PR, ACS; funding acquisition: MEG, MS; investigation: AV, MF, YC, KS; methodology: MF, PR, MEG, MS; project administration: ACS, MEG, MS; resources: MEG, MS; software: AV, MF, PR, MEG; supervision: MEG, MS; validation: AV, MF, YC, PR, KS, ACS, MEG, MS; visualization: AV, MF, YC, ACS, MS; writing – original draft: AV, MF, ACS, MEG, MS; writing – reviewing C editing: AV, MF, YC, ACS, MEG, MS.

## 9 Supplementary Material

### 9.1 Mathematical Model

The mathematical model is a 22-dimensional system of stochastic differential equations designed to describe the behavior of ion channels and their effects on the membrane potential. *I*_LVA_ represents a low-voltage-activated calcium current, while *I*_Na_ and *I*_K_ are the transient sodium and potassium currents responsible for action potential generation. *I*_BK_ is a large-conductance potassium channel that is activated by both a depolarized membrane potential and an increased intracellular calcium concentration, and *I*_L_ is the leak current. Additionally, the Orco channel is a leaky, non-specific ion channel gated by cAMP, with the cAMP concentration oscillating in a circadian rhythm. To model the Orco conductance, we used a sinusoidal function of Zeitgeber time (ZT), independent of the membrane potential, as shown in Figure 2B, C. Each of these ion channels has been reported in vitro in previous studies (Dolzer et al., 2021).

The deterministic part of the 22D model is given by the mean field of the channel-based Langevin model. This approach was introduced by Fox and Lu (1994) for a 14D HH model. In their framework, one dimension tracks the membrane potential, while the other 13 dimensions correspond to the fraction of channels in each of the five states in the potassium (K^+^) Markov chain and the eight states in the sodium (Na^+^) Markov chain. Pu and Thomas (2020) demonstrated that the deterministic 14D model and the original 4D HH model are dynamically equivalent, meaning that each solution of the 4D model corresponds to a solution of the 14D model.

Expanding upon Pu and Thomas’ methodology to represent our system, we added one dimension to track the calcium concentration, six dimensions for each possible state of the LVA channel, and two dimensions for each possible state of the BK channel. The state vectors for each ion channel are therefore defined as follows: a five-component vector N^K^ for the K^+^ gates, an eight-component vector N^Na^ for the Na^+^ gates, a six-component vector N^LVA^ for the LVA gates, and a two-component vector N^BK^ for the BK gates. The state vectors are represented as:

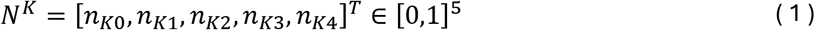

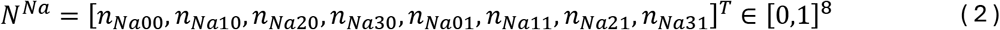

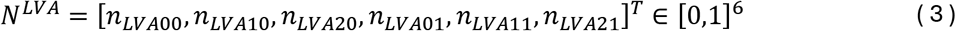

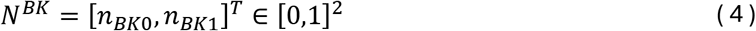

The K^+^ channel consists of four independent n (activation) gates, forming a five-vertex channel-state diagram with eight directed edges. The channel conducts a current only when it is in the rightmost state (Figure 2D). Similarly, the Na^+^ channel is composed of three identical n gates and one h (inactivation) gate, forming an 8-vertex diagram with 20 directed edges, one of which is conducting. The LVA channel includes two identical n gates and one h gate, resulting in a 6-vertex diagram with 14 directed edges, one of which is conducting. The BK channel consists of a single n gate, forming a 2-vertex diagram with two directed edges, one of which is conducting. The Markov chain for every channel is shown in Figure 2B.

For a sufficiently large number of channels, Schmidt and Thomas (2014) and Schmidt et al. (2018) demonstrated that the equations governing the system can be approximated by a multidimensional Langevin equation, as follows:

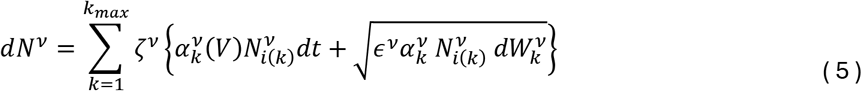

Where the set *ν* contains the four voltage-gated ionic currents included in the model:

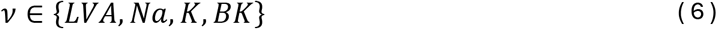

Here *ζ^v^* is the stoichiometry vector for the k^th^ directed edge. If we write i(k) for the source node and j(k) for the destination node of edge k, then *ζ^v^ = e^v^_j(k)_ − e^v^_j(k)_*. The voltage-dependent per capita transition rate along the k^th^ edge is *α^v^_K_(V)* and *N^v^_i(K)_* is the fractional occupancy of the source node for the k^th^ transition. The finite-time increment in the Poisson process is approximated by a gaussian process, namely, the increment *dW^v^_k_* in a Wiener (Brownian motion) process associated with each directed edge. These independent noise terms are scaled by *ϵ = 1/N^v^_Tot_*, where *N^v^_Tot_* represents the total number of that specific ion channel.

The membrane potential is governed by the current balance equation:

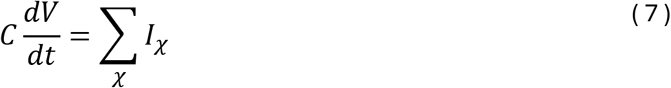

where the set *χ* contains the six ionic currents included in the model:

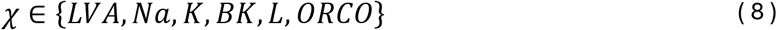

Each of the six ionic currents included in the model are described as:

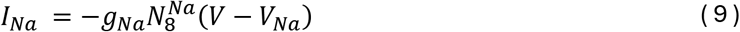

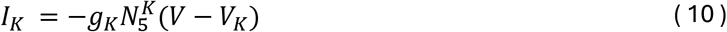

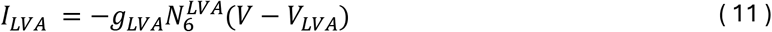

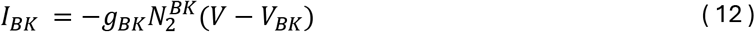

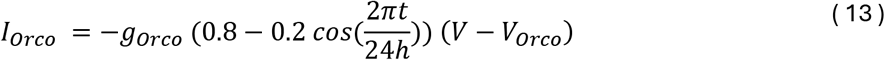

Here, *g_χ_* is the maximal conductance, *V* is the membrane potential, and *V_χ_* is the reversal potential for each ion current. The parameter values for each current can be found in Table 1.

**Table 1:** Parameter values for model currents.

| Current | $g_x(mS/cm^2)$ | $V_x(mV)$ | $N_{Tot}^v$ |
| --- | --- | --- | --- |
| $I_L$ | 4.3 | -62.5 | - |
| $I_{LVA}$ | 10.0 | 120 | 1800 |
| $I_{NA}$ | 29.17 | 45.0 | 6000 |
| $I_K$ | 12.96 | -105.0 | 1800 |
| $I_{BK}$ | 9.0 | -105.0 | 1800 |
| $I_{ORCO}$ | 5.0 | 0.0 | - |

Each transition rate *α^v^_k_(V)* is calculated in the following way:

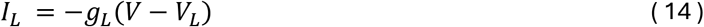

where *p^v^_j(k)∞_(V)* is the steady state value of the specific gate that switches states in the destination state and *τ_p_*(*V*) is the time constant of said gate. The steady state activation values and time constants for the LVA, Na, K, and BK channels were based on Viertel and Borisyuk (2019). The steady state functions for the activation gates are denoted with *m^v^_∞_*, and those for the inactivation gates with *h^v^_∞_*.

Each gate’s steady state activation function, except for the BK channel, takes the form:

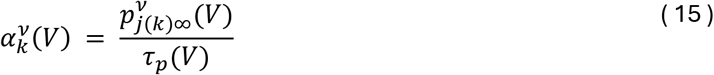

The *I*_BK_ current is dependent on the intracellular calcium concentration and membrane potential as represented by the following equations:

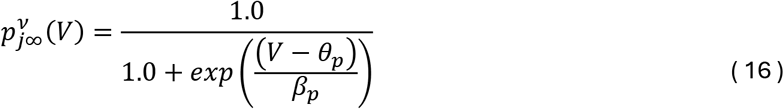

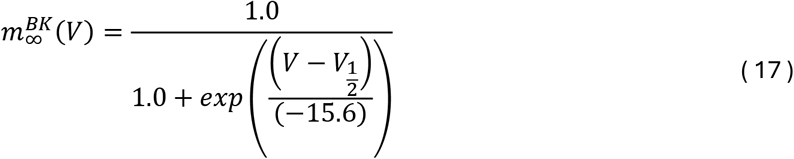

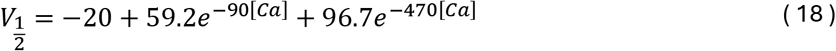

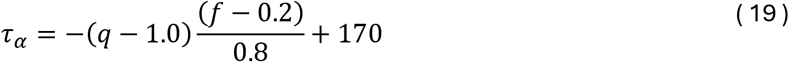

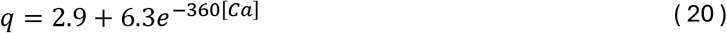

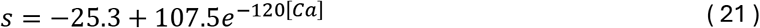

The intracellular concentration of calcium ions is described by the following equation:

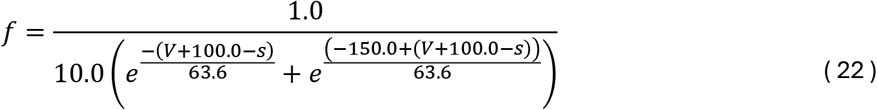

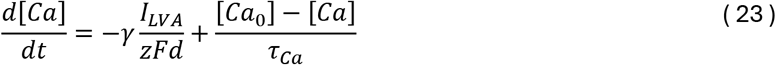

where *Ca*_0_ is the steady state calcium concentration, *γ* describes calcium buffering, *z* is the ionic valence of calcium, *F* is Faraday’s constant and *d* is the depth in microns at which calcium ions concentrate near the membrane.

The parameters of each steady state activation function as well as the time constant for each gating value can be found in Table 2. Values of all membrane and calcium dynamics parameters are listed in Table 3.

**Table 2:** Gating variable parameter values.

| Variable | $\theta$ | $\beta$ | $\tau(ms)$ |
| --- | --- | --- | --- |
| $m^{LVA}$ | -30 | -4.8916 | $\frac{40.0}{\cosh\left(\frac{(V + 37.1)}{9.7832}\right)}$ |
| $h^{LVA}$ | -59.2 | 11.2326 | $\frac{350.0}{\cosh\left(\frac{(V + 59.2)}{22.4652}\right)}$ |
| $m^{Na}$ | -25.0 | -6.5 | 0.2 |
| $h^{Na}$ | -26.0 | 9.0 | $\frac{10.0}{\cosh\left(\frac{(V + 26.0)}{18.0}\right)}$ |
| $m^K$ | -26.0 | -9.0 | $\frac{10.0}{\cosh\left(\frac{(V + 26.0)}{18.0}\right)}$ |

**Table 3:** Membrane and calcium dynamics parameters.

| Parameter | Value | Unit |
| --- | --- | --- |
| C | 21.0 | $\mu F/cm^2$ |
| [Ca<sub>0</sub>] | $2 \times 10^{-5}$ | mM |
| $\tau_{Ca}$ | 80.0 | ms |
| $\gamma$ | 1.0 | - |
| d | 1.0 | $\mu m$ |

The stochastic differential equations were integrated numerically using the Euler–Maruyama method with a fixed time step of Δt = 0.008 ms. Two separate simulations were performed. For Figures 8C and 8D, a long-term simulation spanning 3 days was carried out with the Orco current following the circadian modulation given by Equation (13). For Figure 8B, a separate 1-hour simulation was performed with the Orco conductance held fixed at gOrco = 7.8 mS/cm², corresponding to a high-conductance, high-activity condition.

---

**Figure 3 – Supplement:**
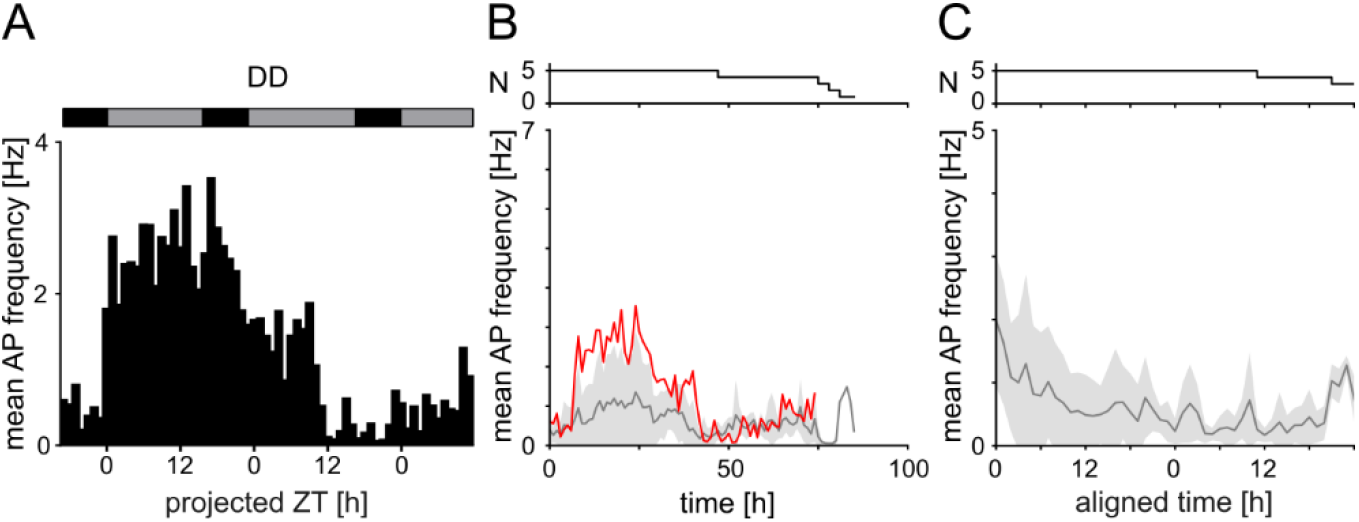
Spontaneous spiking activity of ORNs of pheromone-exposed male *M. sexta*. To enhance synchronization, males were exposed to females and pheromones as additional Zeitgeber for 30 mins at ZT 16 before the start of the experiment. Recordings were obtained over multiple days in DD. **(A)** The spontaneous spike frequency of one long-term ORN recording in constant darkness (DD, the expected times of lights on are represented by grey bars above) **(B)** Mean spontaneous ORN spike frequency across all DD animals in 1 h bins. Data from the individual in **A** is highlighted in red. **(C)** Mean spontaneous ORN spike frequency across all DD animals in 1 h bins during the first 48 hours after aligning the time to the first maximum in spontaneous activity (see Methods). No circadian activity was evident in these pheromone-exposed males.

